# Molecular mechanisms of tubulogenesis revealed in the sea star hydro-vascular organ

**DOI:** 10.1101/2022.08.25.505020

**Authors:** Margherita Perillo, S. Zachary Swartz, Cosmo Pieplow, Gary M. Wessel

## Abstract

A fundamental goal in the organogenesis field is to understand how cells organize into tubular shapes. Toward this aim, we have established the hydro-vascular organ in the sea star *Patiria miniata* as a model for tubulogenesis. In this animal, bilateral tubes grow out from the tip of the developing gut, and precisely extend to specific sites in the larva. This growth requires cell migration coupled with mitosis in distinct zones. Cell proliferation requires FGF signaling, whereas the three-dimensional orientation of the organ depends on Wnt signaling. Specification and maintenance of tube cell fate requires Delta/Notch signaling. Moreover, we identify target genes of the FGF pathway that contribute to tube morphology, revealing molecular mechanisms for tube outgrowth. Finally, we report that FGF activates the Six1/2 transcription factor, which serves as an evolutionarily ancient regulator of branching morphogenesis. This study uncovers novel mechanisms of tubulogenesis in vivo and we propose that cellular dynamics in the sea star hydro-vascular organ represents a key comparison for understanding the evolution of vertebrate organs.

**Highlights:**

❖ The hydro-vascular organ of the sea star presents a valuable model of tubulogenesis
❖ In this organ tube extension is driven by cell migration coupled with cell proliferation at specific growth zones
❖ The Wnt pathway controls directional outgrowth
❖ The FGF pathway promotes regionalized cell proliferation
❖ The Notch/Delta pathway is essential in cell fate repression in tubulogenesis
❖ A screen of FGF function revealed essential target gene expression, including the transcription factor Six1/2
❖ Within a sister group to chordates, the sea star will reveal ancient mechanisms of tubulogenesis

## Introduction

The coordinated organization of cells into a precise three-dimensional architecture is critical for establishing functional organs in development. Tubulogenesis, the process by which hollow tubes form, is an important transient step in the development of many organs, including the neural tube, heart and pancreas (Iruela-Arispe & Beitel, 2013). In addition, many vital organs are ultimately organized into tubular shapes with the essential scope of transporting fluids, gasses, or cells, as in the case of kidneys, mammary glands, and vasculature (Lubarsky & Krasnow, 2003).

A great diversity of mechanisms are used to form hollow tubes in animals, including how cells reach their final position and distinct external cues that guide the proper organ shape (Iruela-Arispe & Beitel, 2013; Lubarsky & Krasnow, 2003). Abnormalities in these processes can cause congenital disorders, dysfunctional or displaced organs, and loss of organ symmetry. Yet, because of the great diversity of tubular structures, the general and conserved mechanisms of tube formation in organogenesis are not completely understood (Iruela-Arispe & Beitel, 2013; Lubarsky & Krasnow, 2003; Sigurbjörnsdóttir et al., 2014).

Our knowledge of tube formation is largely derived from specific organs within selected model systems, including *D. melanogaster*, C. *elegans*, C. *intestinalis*, zebrafish, mice, cell culture, and organoids (Buechner et al., 2020; Denker & Di, 2012; Hirashima et al., 2017; Kerman et al., 2006; Kim et al., 2021; Mailleux et al., 2008; Riccio et al., 2016; Vasilyev et al., 2012; Warburton et al., 2000). However, there are some key differences amongst these systems. For example, while it is broadly true that cells within tubular epithelia must become polarized, an essential step for lumen formation, polarization is acquired differently in diverse organs and species (Best, 2019; Buechner et al., 2020). Similarly, cell migration, morphogenetic movements, and mitotic division are important for forming the three-dimensional structures. However, each of these parameters contribute to different extents in different species. For instance, while in *Drosophila* tubular organs cells undergo massive proliferation before migration, in vertebrates cell proliferation and migration are coupled (Denholm, 2013; Hogan & Kolodziej, 2002; Kerman et al., 2006; Kerman et al., 2008; Mollard & Dziadek, 1998; Samakovlis et al., 1996; Skaer, 1989; Vasilyev et al., 2012; Weaver et al., 2000). From an evolutionary perspective, it is therefore an important challenge to determine whether these different cell mechanics in organogenesis reflect the evolution of organs, and whether a basal tubulogenesis toolkit may exist.

Similarly, it remains unclear whether there exists a core and conserved network of signaling pathways that guide cell movements and fate decisions in tubulogenesis. For example, in angiogenesis and vasculogenesis, Notch signaling regulates the differentiation of vascular endothelial cells (David Morrow et al., 2008; Sainson et al., 2005; Zhong et al., 2001). However, whether Notch participates in tube formation in other contexts remains poorly understood. In addition, Wnt signaling controls the development of branching organs with complex architectures in both vertebrates and invertebrates (Chihara & Hayashi, 2000; De Langhe et al., 2005; Karner et al., 2009; Li et al., 2002; Llimargas & Lawrence, 2001; Loscertales et al., 2008; Matsuyama et al., 2009; Moura et al., 2014; Mucenski et al., 2003; Saburi et al., 2008; Yu et al., 2009). However, because of the complexity of these organs, other morphological aspects besides branching have been difficult to investigate. Furthermore, FGF signaling is important for tubulogenesis across species, including the mouse male ureteric bud, lungs and kidney (Lebeche et al., 1999; Litwin et al., 2015; Ohuchi et al., 2000; Okazawa et al., 2015; Park et al., 1998; Peters et al., 1994; Qiao et al., 1999; Zhao et al., 2004).However, many critical aspects of FGF action are still poorly understood, including the downstream targets of FGF signaling, and the conservation of these target genes across organ types and species.

The canonical model systems from which most of our knowledge of tubulogenesis derives generally belong to two main phylogenetic groups: ecdysozoans (e.g. worms and flies) and chordates (e.g. tunicates and vertebrates) (Figure 1 A). Echinoderms, which include sea urchins and sea stars, are part of the sister group to chordates and as such occupy a critical phylogenetic position to understand evolution of vertebrate organs, while offering a series of experimental advantages (Annunziata et al., 2019; Annunziata et al., 2013; Annunziata et al., 2014; Arnone et al., 2016; Ben-Tabou de-Leon, 2022; Cary et al., 2019; Cary & Hinman, 2017; Cheatle Jarvela & Hinman, 2014; Hinman & Cheatle Jarvela, 2014; Meyer & Hinman, 2022; Morgulis et al., 2019; Morgulis et al., 2021; Tarsis et al., 2022; Wolff & Hinman, 2021). Here, we use the sea star *Patiria miniata* as a powerful system to study tubulogenesis within an intact organism. The sea star larva is genetically tractable, optically transparent, and forms a tubular coelom organ called the anterior coeloms, coelomic sacs, or the hydro-axocoel (**hydro-vascular organ** from here on) (Child, 1941; Horstadius, 1973; Hyman, 1955). The hydro-vascular organ functions as a hydrostatic skeleton to allow the larva to balance in the water column, and as such it is a simple, but vital organ (Newton & Potts, 1993; Potts, 2003). Using live imaging and functional approaches, we identify the intrinsic (cell migration and proliferation) and extrinsic (external cues) factors driving the first steps of tubulogenesis in this novel system. We reveal that in an early branching deuterostome, directional cell migration coupled with cell division drives tubulogenesis. The initial tube outgrowth, the oriented elongation, and the differentiation of mesodermal cells as epithelial tube cells rely of the FGF, Wnt and Delta/Notch pathways. Our work defines a basic toolkit from where the chordate tubular organs might have evolved.

**Figure 1.**
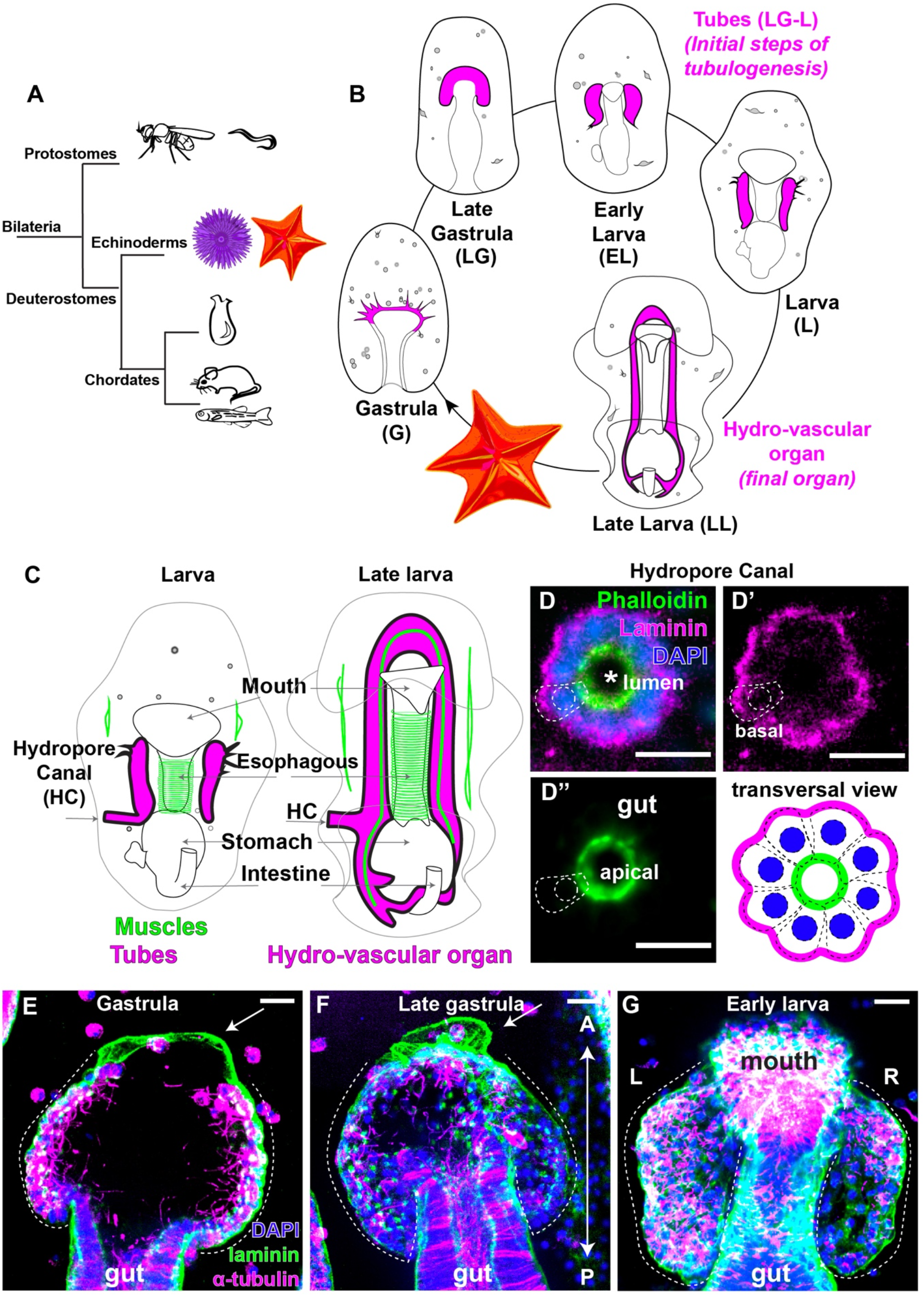
Development of the sea star larva hydro-vascular organ. **A)** Phylogenetic relationships of the main bilaterian groups. **B)** Summary of sea star larval development with a focus on the hydro-vascular organ (in magenta). The hydro-vascular organ comes from mesodermal precursors located at the tip of the growing gut in gastrula (G). Tubulogenesis starts in the late gastrula (LG), continues in the early larva (EL) and larval stages (L1 to L3). In larval stages, the initial tubes that form the organ elongate posteriorly. The left tube forms the hydropore canal, an opening toward the outside environment. The organ is fully grown in late larva (LL), when the left and right tubes merge to form a closed system. **C)** Summary of the main larval tissues and organs. **D-D”)** Transversal view of the tube trough the hydropore canal showing that the epithelium is polarized with actin on the apical side of the cells (facing the lumen) and laminin on the cell basal side. Dotted lines indicate the polarity of a single tube cell. Scale bars 10 μm. **E-G)** Laminin staining showing the basal lamina protrusion (arrow) in gastrula and late gastrula stages that redistributes in later larval stages with the progression of tube morphogenesis. Alpha tubulin marks cilia in the tube lumen (insert). Scale bars = 20 μm. L, left; R, right.

## Results

### The hydro-vascular organ acquires apical-basal polarity early in its development

Tubulogenesis is a highly dynamic process of individualized cell movements and tissue-level morphogenesis. To define the nature of sea star tube formation, we took advantage of the optical clarity of the larvae and developed techniques to immobilize and live image morphogenesis continuously from gastrulation to early larval stages (Figure 1- Video 1). To standardize the different steps of tubulogenesis, we defined a naming scheme to summarize the critical stages of tube development (Figure 1B, Figure 1 -Table 1). The hydro-vascular organ derives from mesodermal progenitors (Yankura et al., 2010) that, from the tip of the growing gut, divide into two bilateral tubes and grow towards the posterior end of the larva (Figure 1- Video 2). The two tubes ultimately merge to form a contiguous system with one opening towards the external environment, the hydropore canal (Figure 1 C, D-D”). This entire process happens in approximatively 15 hours, as shown in the video (Figure 1 -Video 1).

Our first goal was to define the tissue morphologically. A fundamental feature of tubular organs is the presence of a polarized epithelium, and to determine this tissue morphology, we tested whether the cells displayed apical-basal polarity by visualizing actin and the basal lamina component laminin. A transverse view through the hydropore canal showed that the tube epithelium is polarized, with the apical side of the cells rich in actin cytoskeleton facing the lumen and a basal lamina delimiting the blastocoel environment (Figure 1 D-D”). We found that the tubes were continuously enwrapped by a layer of laminin from where they originated (Figure 1 E, F, G). At the onset of tubulogenesis (gastrula stage), before the two tubes fully separated from the gut, cells were already polarized, as shown by the presence of cilia that defines the apical surface on the luminal side (Figure 1 E, Figure 1 -Figure supplement 1). In the fully formed tubes we detected cilia beating (Figure 1 -Video 3) suggesting that fluids are circulating inside the lumen of the tubes and that these are not primary cilia. Although the basal lamina was generally contiguous with the tube epithelium, we observed a laminin protrusion on the anterior side of the tube that was devoid of cells (Figure 1 E, F arrows), suggesting that cells left the laminin network on the anterior side after their migration toward the posterior end. These results indicate that the progenitors of the hydro-vascular organ are polarized early in the development of this structure and form a dynamic epithelium.

### Tubulogenesis is driven by cell migration along the posterior and medial axes

By live imaging, we determined that the tubes elongate from cells at the top of the growing gut (Figure 1- Video 1). We tracked single cell movements from late gastrula to early larva stage in embryos expressing H2B-GFP to label nuclei. We recorded cell movements with respect to the anterior-posterior axis (A-P) and the left-right axis (L-R) (Figure 2 -Videos 1-4 and Figure 2 -Figure supplements 1 and 2). Strikingly, we identified two distinct phases of active cell migration. In the first phase, cells made substantial movements towards the posterior end of the embryo (Figure 2 A-B, E-G). On average, cells made the most progress along the A-P axis and completed most of their overall movements during this first phase, calculated as the total migration (Figure 2H). In the second phase, cells maintained their relative positions along the A-P axis (Figure 2 C-D, E-G) and moved on the L-R axis more than on the A-P axis (Figure 2 H). We thereby uncovered a two-phase program of cell movements, in which the cells first move towards the posterior axis to promote rapid tube elongation, and then along the L-R axis to promote tube expansion.

**Figure 2.**
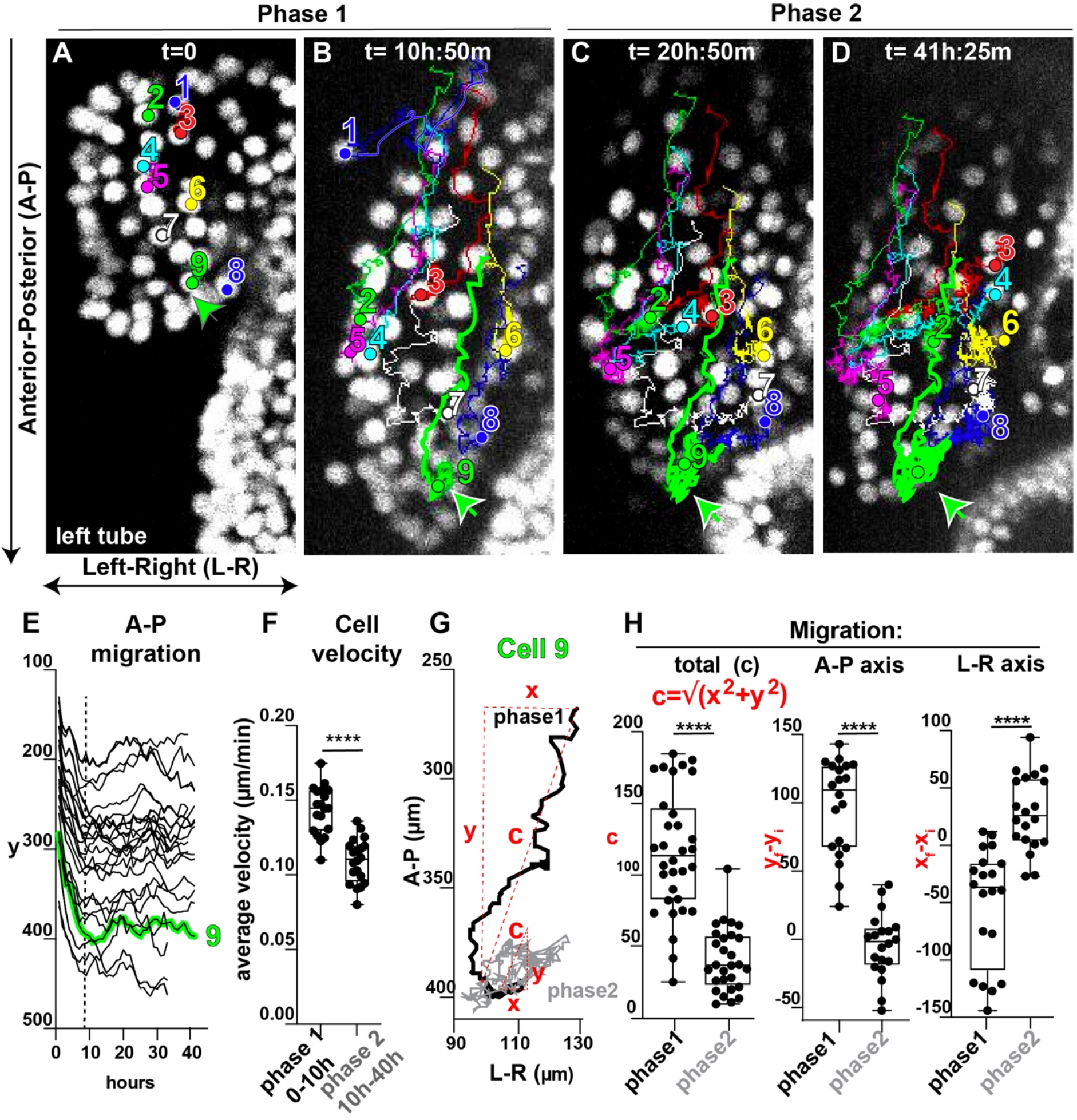
Cell migration guides tube extension and expansion. **A-D)** Time lapse images of the tubes expressing H2B-GFP to track movements of single nuclei. Tracked cells are labeled with unique colors and numbers to visualize trajectories. In 2D projections, cell movements are recorded along the A-P axis (advancing direction) and the L-R axis. Note that in A and B, cell 1 has undergone an epithelial to mesenchymal transition (EMT) and migrates into the blastocoel, away from the elongating tube. Still images are from Figure 2 -Video 2. **E)** Cell migration along the A-P axis of the embryo (the y axis of the movie). Cell movements along the A-P axis are represented relative to hours. Cell 9 from A-D) is highlighted in green to show the two stages of migration. Around 8 hours into the movies, cells stop their rapid migration toward the posterior end and remain relatively still with respect to the A-P axis. Graph includes cell trajectories of both tubes for 2 independent movies (n=30 cells). **F)** The trajectory of Cell 9 exemplifies cell movement along A-P and L-R axes. After a rapid migration towards the posterior end, cells move along the L-R axis. C represents the total cell displacement and is calculated as shown. N= 39 cells from two independent movies. **G)** Average cell velocity for phase 1 and 2 for 2 independent movies. **H)** In phase 1, the maximum cell displacement occurs along the A-P axis. In phase 2, cell displacement happens along the L-R axis. Track data for n= 50 cells from two independent movies. For Student’s t-test **** p-value is < 0.0001.

### Mitotic growth zones contribute to tube elongation

We next tested whether mitosis contributes to construction of the hydro-vascular organ. Indeed, we observed numerous mitotic events during both stages of tube cell migration (example in Figure 3 -Figure supplement 1A and Figure 2 -Videos). We therefore quantified the relative frequency of mitosis along the tubes using EdU pulse experiments. The overall cell proliferation along the tube increased from LG until L1 (first larval stage) and then sharply decreased until the L3 (last larval stage) (Figure 3A). We next asked whether cell proliferation occurs stochastically along the tube, or in spatially localized growth zones. We therefore quantified EdU positive cells within three regions of the tube: the stalk (anterior, where cell migration starts, close to the growing gut), the middle, and tip cells (posterior-most) (Figure 3 B-D and Figure 3- Figure supplement 1 B-E). In the initial stages of growth, late gastrula and early larva, cell proliferation was higher in the middle and tip cells than the stalk cells (Figure 3 E). In the first larva stages (L1-L2), the middle and tip zone cells were dividing the most, whereas by L3 stage, proliferation had decreased throughout the tube (Figure 3 D, E).

**Figure 3.**
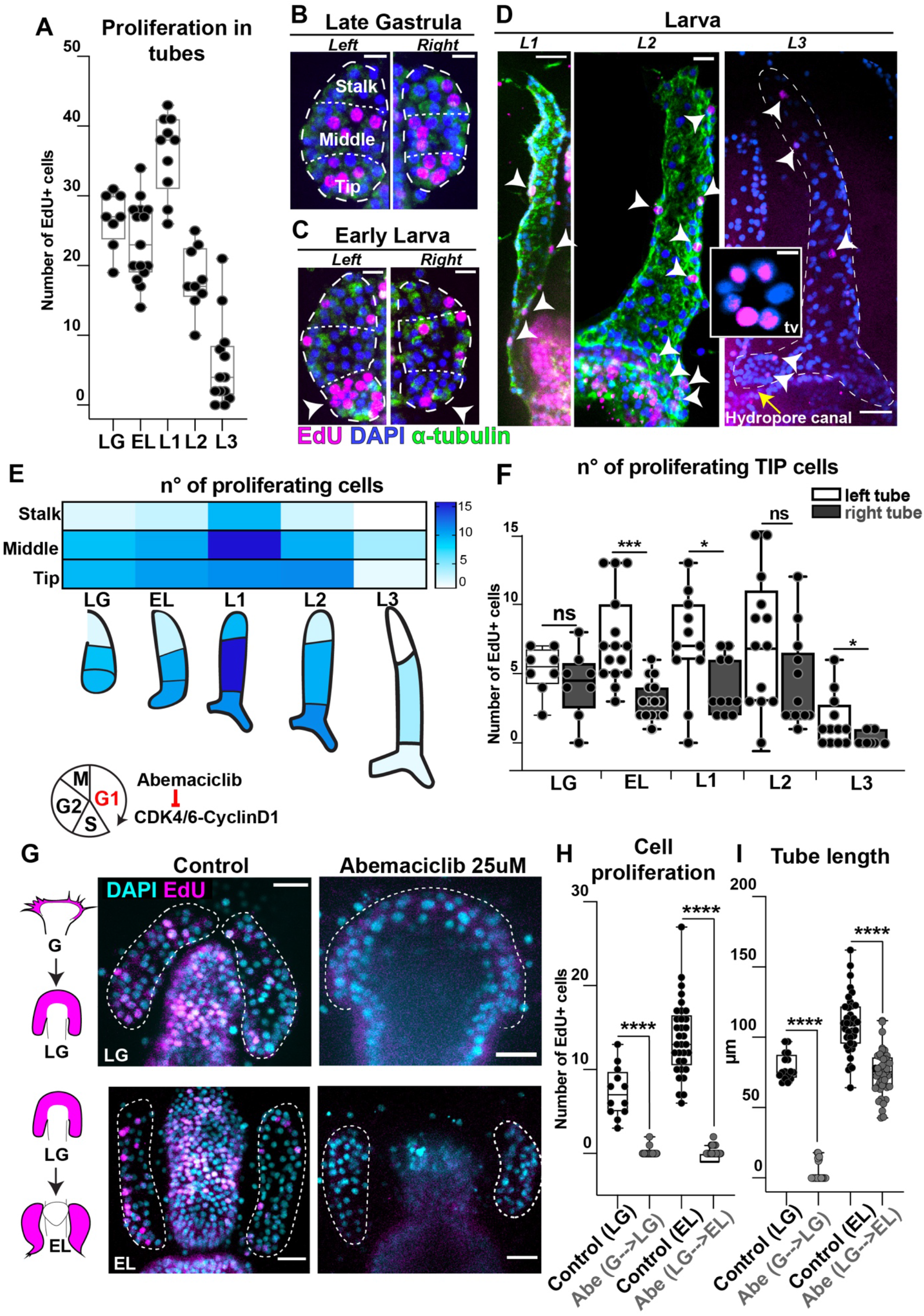
Cell division during tube outgrowth. **A)** EdU pulse experiments to measure frequency of cell proliferation during tubulogenesis from tube outgrowth until larva stages (as defined in Figure 1- Table 1). Number of total larvae analyzed n=60. Number of Edu +cells are plotted by stages. **B, C)** EdU staining (magenta) in the three areas of the tube: stalk, middle and tip cells. Arrowheads in C indicate more cell proliferation in the tip of the left tube than the right one. Scale bar 10 μm **D)** EdU staining in larval stages. Arrowheads indicate nuclei with incorporated EdU. Insert shows a transverse view (tv) of the hydropore canal from another specimen with nuclei stained with EdU. Scale bar 20 μm, insert 5 μm. **E)** Heatmap showing frequency of cell proliferation in the three tube areas during tubulogenesis (white is zero, dark blue is max). **F)** Frequency of cell proliferation in the tip cells for left and right tubes. N= 60 total number of larvae. **G)** EdU staining in larvae treated with Abemaciclib to arrest cells in G1 from gastrula to late gastrula (during tube outgrowth) or from late gastrula to early larva (during tube elongation). Dotted lines indicate the tubes. Scale bar 20 μm. **H)** EdU incorporation (number of EdU + cells) in presence of Abemaciclib to test inhibition in cell cycle and **(I)** effects on tube outgrowth and elongation. Number of larvae n=45 for controls and n= 66 for treated. For Student’s t-test ****p-value is < 0.0001. Ns= not significant. LG (late gastrula); EL (early larva); L1 (larva stage 1); L2 (larva stage 2); L3 (larva stage 3).

The left and right tubes are morphologically distinct, with the left tube specifically branching out to create the hydropore canal, and so we tested whether there were proliferative differences between them. We found that the tip cells of the left tube proliferated more than the right one, from the EL to L3 stages (Figure 3 F). Cell proliferation was particularly active in the hydropore canal (Insert in Figure 3 D, transversal view). We found no difference in cell proliferation at the middle zone between left and right tubes (Figure 3 -Figure supplement 1F). In the late larva, cell proliferation occurred exclusively in the posterior-most region of the final tube organ (Figure 3- Figure supplement 1G). In summary, an extensive cell division during tubulogenesis occurs within specific zones of the middle and tip areas of the tubes, and especially within the left tube tip.

We next asked whether cell division was required for tubulogenesis by blocking cell cycle progression in G1 using the Cdk4/6 inhibitor Abemaciclib (LY2835219) (Palumbo et al., 2019). First, to test whether cell proliferation was necessary for initial tube outgrowth, we added the inhibitor from the end of gastrulation until the early larva stage. Importantly, and consistent with a G1 arrest and failure to replicate DNA, the tube cells did not incorporate EdU under these conditions (Figure 3 G, H). Strikingly, when cell cycle progression was blocked, tube outgrowth was fully prevented (Figure 3 G). Second, to test whether ongoing cell division was needed after initial outgrowth to support continued elongation, we instead treated embryos from late gastrula to early larva. We found that cell cycle arrest at this stage prevented further elongation, with the length of the tubes remaining the same as when the drug was added (Figure 3 I). These results indicate that cell proliferation is a critical driving force to both initiate and sustain tubulogenesis.

### Delta-Notch signaling provides cell-fate decisions in the tubes

Having defined cell migration and cell division as essential intrinsic forces for tube elongation, we next tested the contribution of extrinsic signaling pathways. In the sea star embryo, Delta-Notch signaling is critical during gastrulation for specifying all mesodermal cell types, including the hydro-vascular organ precursor cells, migratory mesoderm, and muscles (Cary et al., 2020; Hinman & Davidson, 2007). To test whether Delta-Notch signaling could also guide later steps of tube formation we first examined the spatial expression of the Delta ligand after gastrulation and detected Delta mRNA selectively in most tube cells (Figure 4 A). The Notch receptor was instead reported to be ubiquitously expressed (Cary et al., 2020). To define the contribution of the Delta+ tube cells to tubulogenesis we generated Delta knockout embryos. In Delta KO late gastrulae, the tube was not specified and there was an increase of mesodermal cells expressing Erg and Ets1/2 (Figure 4 B-D), markers of mesenchyme cells (Cary et al., 2020; Hinman & Davidson, 2007). These results are consistent with previously reported pharmacological Notch inhibition (Cary et al., 2020).

**Figure 4.**
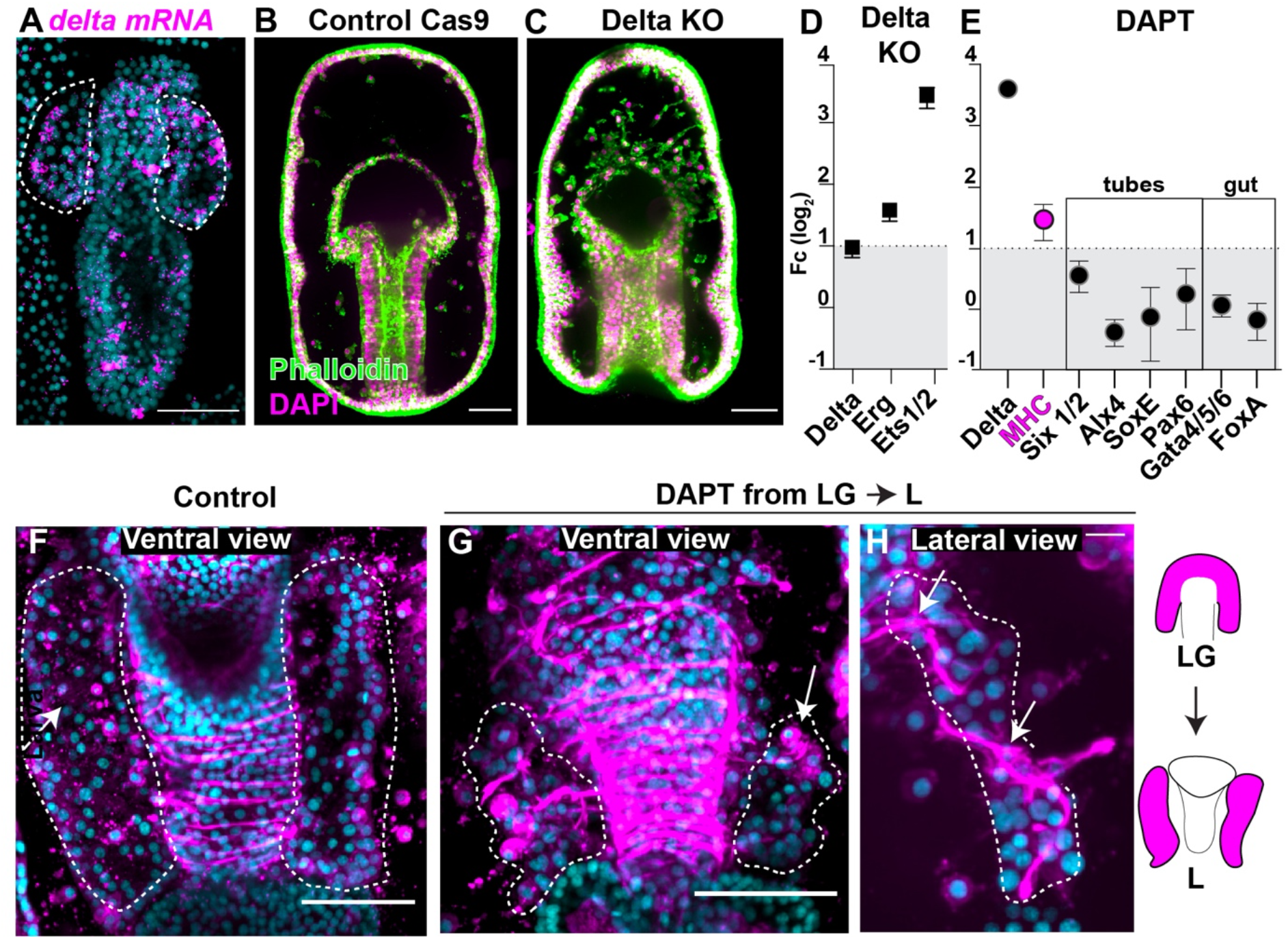
Delta-Notch signaling represses a muscle phenotype in the tubes. **A)** Fluorescent in situ hybridization (FISH) shows Delta gene expression in scattered cells of the gut and the tubes (highlighted by dotted lines). **B, C)** Delta knock out embryos fail to form the epithelial tubes and instead become excess mesenchymal cells. **D)** qPCR showing increase of mesodermal markers Erg, Ets1/2 and Pax6 in delta KO embryos. **E)** The increase in muscle-like cells is also reflected by increase in expression of the muscle marker MHC, while other known marker genes for tubes and the digestive system do not change. **F-H)** Magnification of the tubes from the same larvae shown in Figure 4 -Figure supplement 1. Larvae stained with phalloidin to mark muscles (magenta); Delta-Notch inhibition leads to smaller tubes constricted by muscle fibers that wrap around the tubes (arrows indicate muscles) and the foreguts. Arrows in G show that tube cells become muscle-like. All experiments are representative of 3 biological replicates. A, B, C, F, G scale bar = 50 μm. H scale bar is 10 μm.

To test the contribution of Delta+ cells to later stages of tubulogenesis, we used the γ-secretase inhibitor DAPT to block Delta/Notch signaling with temporal control (Cary et al., 2020; Perillo et al., 2016). When we treated embryos at the end of gastrulation with DAPT, after tube cells were already specified, larvae had smaller tubes (Figure 4 F-H, Figure 4 -Figure supplement 1). Surprisingly, while control tubes did not develop muscles by this stage, Delta-Notch inhibited larvae displayed increased muscle fibers wrapped around the tubes, even resulting in increased constriction (Figure 4 G, H). These muscle fibers appeared to originate from the tube itself (Figure 4 G, H arrows). In further support of excess muscle specification in the absence of Delta/Notch signaling, we detected increased expression of the muscle marker myosin heavy chain (MHC) (Figure 4 E). These results suggest that continued Delta/Notch signaling is important to restrict excess muscle specification.

We next asked how gene expression in the tubes was affected under Delta/Notch inhibition. We tested a panel of known genes expressed in the tubes in this animal, including Six1/2, SoxE, Alx4 (Alx1l) and Pax6 (Cary et al., 2020; Fresques et al., 2014; Khor & Ettensohn, 2020; McCauley et al., 2012; Yankura et al., 2010). We found that the expression of these genes was unchanged (Figure 4 E).

Some Delta+ cells in the digestive system expressed the genes FoxA and GataE (Hinman & Davidson, 2007), which we also found to be unaffected. As seen in previous studies (Cary et al., 2020; Hinman & Davidson, 2007), Delta mRNA expression was increased when Delta/Notch signaling is blocked, indicating that the perturbation was effective (Figure 4 E). These data suggest that although early Delta/Notch signaling is required for tube cell specification (Cary et al., 2020; Hinman & Davidson, 2007), known markers of the tubes were not affected by later stage Notch inhibition. Instead, we found that Notch signaling at later stages prevents the uncontrolled specification of muscle-like cells from the tube cells.

### Wnt signaling guides directional growth of the tubes

Since the sea star tubes stereotypically extend along the anterior-posterior axis, we asked what signals guide this polarized growth. The Wnt pathway is a broadly conserved regulator of polarity at the organismal level and branching for many tubular organs (De Langhe et al., 2005; Li et al., 2002; Llimargas & Lawrence, 2001; Loscertales et al., 2008; Moura et al., 2014; Mucenski et al., 2003) and inhibition of Wnt signaling in the early stages of sea star development blocks the formation of endomesodermal structures (McCauley et al., 2015; Swartz et al., 2021). To test the later contribution of Wnt signaling to tubulogenesis, we blocked secretion of all Wnt ligands after gastrulation with the porcupine inhibitor ETC-159 (Katoh, 2017). Importantly, treatment at this stage does not grossly affect primary body axis or endomesoderm specification, which are established earlier in development. We found that in Wnt-inhibited larvae, the tubes elongated towards the dorsal ectoderm instead of posteriorly, as in controls (Figure 5 A, B). As a control, we tested the expression of a known beta catenin target, Wnt8 (McCauley et al., 2015), and found it to be downregulated with ETC-159 treatment (Figure 5- Figure supplement 1). However, despite the loss of tube directional extension, the expression of tube and muscle associated genes were unchanged (Figure 5- Figure supplement 1). These data suggests that Wnt signaling drives directional growth of the tubes towards the posterior end of the larva but is dispensable for the sustained expression of tube associated genes.

**Figure 5.**
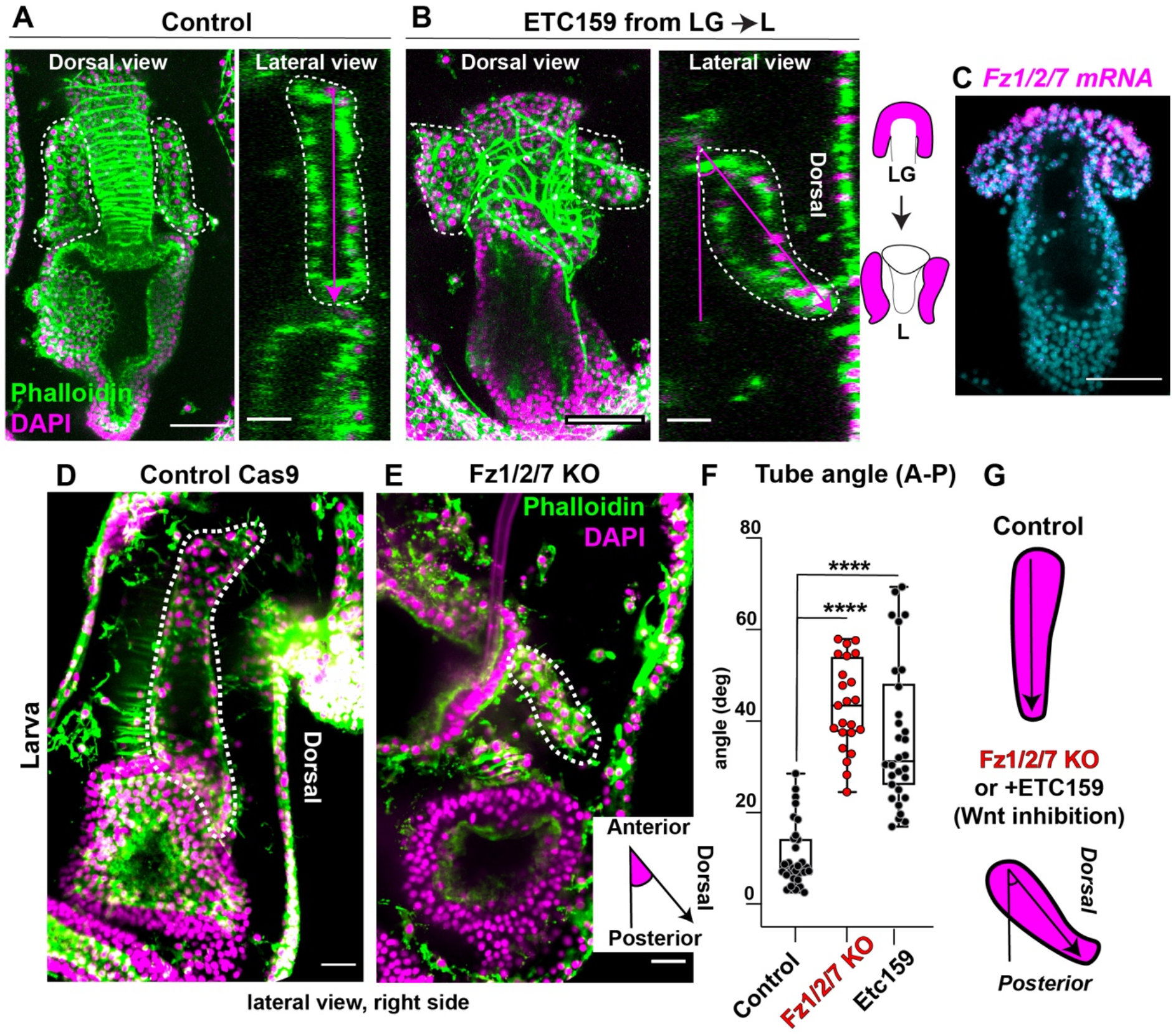
Wnt signaling directs tube orientation through the Fzd1/2/7 receptor. **A, B)** Wnt inhibition with ETC159 from late gastrula to L2 caused a change in orientation of the tube outgrowth from anterior-posterior to anterior-dorsal. Confocal z-stacks are dorsal views and lateral views obtained from orthogonal projections for the same larvae. **C)** FISH showing that the Wnt receptor Fzd1/2/7 is expressed in the tube cells in early larvae. **D**, **E)** In control larvae tubes grow parallel to the anterior-posterior axis, while in Fzd1/2/7 knock out larvae the tubes grow towards the dorsal side and recapitulate the same tube orientation defects seen with ETC159 treatment. **F)** Tube orientation measured as the angle that the tube makes with respect to the anterior-posterior axis of the larva. Number of tubes measured is n= 30 for controls, n=30 for ETC159 treated and n=23 for Fzd1/2/7 KO. **G)** Schematic summarizes the observed phenotypes. For Student’s t-test ****p-value is < 0.0001. All experiments are representative of 3 biological replicates. A, B, D, E, F scale bar is 50 μm. A and B lateral views are 20 μm.

We next sought to define the specific Wnt signaling pathway and its receptor used in tubulogenesis. A prior study identified Fz1/2/7 as initially localized broadly in the tip of the archenteron, and later selectively in the tubes that form the hydro-vascular organ (McCauley et al., 2013). We further defined the localization of Fz1/2/7 mRNA, and found its expression enriched in the anterior tube cells (Figure 5 C). To test whether tube orientation was dependent on Fz1/2/7, we generated Fz1/2/7 knockout embryos and found that the tubes elongated dorsally instead of posteriorly, which phenocopied Wnt inhibition by EC-159 (Figure 5 D, E). To assess the directionality of tube elongation we measured the angle that each tube formed with the larval anterior-posterior axis and found that this angle is similar in Fz1/2/7 KO and ETC-159 treated embryos but greater than in controls (Figure 5 F-G). We conclude that Wnt signaling via the Fz1/2/7 receptor guides oriented growth of the left and right tubes towards the posterior end of the larva, the site where the two tubes will eventually fuse to form the hydro-vascular organ of the late larva.

### FGF signaling initiates tube outgrowth and elongation through pERK by controlling cell proliferation

FGF signaling has been implicated in the morphogenesis of many kinds of tubular structures (Andrew & Ewald, 2010; Sutherland et al., 1996). Consistent with a potential role in forming the tubes, we found that the FGF receptor is expressed by scattered tube and esophageal cells (Figure 6 A). To test whether FGF signaling is required for tube formation, we used the FGFR inhibitor SU540 (Andrikou et al., 2015; Czarkwiani et al., 2021; Perillo et al., 2016). Strikingly, we found that larvae in which FGFR was inhibited following gastrulation had partially missing tubes (Figure 6 B, C). As a second approach, we used the inhibitor U0126 to block phosphorylation of the FGFR downstream effector kinase MEK (Fernandez-Serra et al., 2004). This treatment also resulted in a partial lack of tubes, similar to the upstream FGFR inhibition (Figure 6 D). We then knocked out FGFR and found that also with this third approach the tubes developed only partially (Figure 6 E, cartoon in F), with a significant decrease in tube length and area compared to control early larvae (Figure 6 G, H). Strikingly, aside from lacking the tubes, the overall anatomy of the late larva was normal (Figure 6- Figure supplement 1), indicating a highly selective role of FGF signaling in tubulogenesis. Consistent with inactivation of the FGF pathway, we found that all of these manipulations resulted in a decrease in nuclear pERK, the target of MEK (Figure 6 J, K, Figure 6- Figure supplement 2 A, B). Moreover, when FGFR was knocked out or if its kinase domain was blocked, genes normally expressed in the tubes and the muscle marker MHC were significantly downregulated (Figure 6 I). After two weeks, while controls were alive the FGFR KO larvae died, suggesting that lack of the tube organ is critical for larval survival.

**Figure 6.**
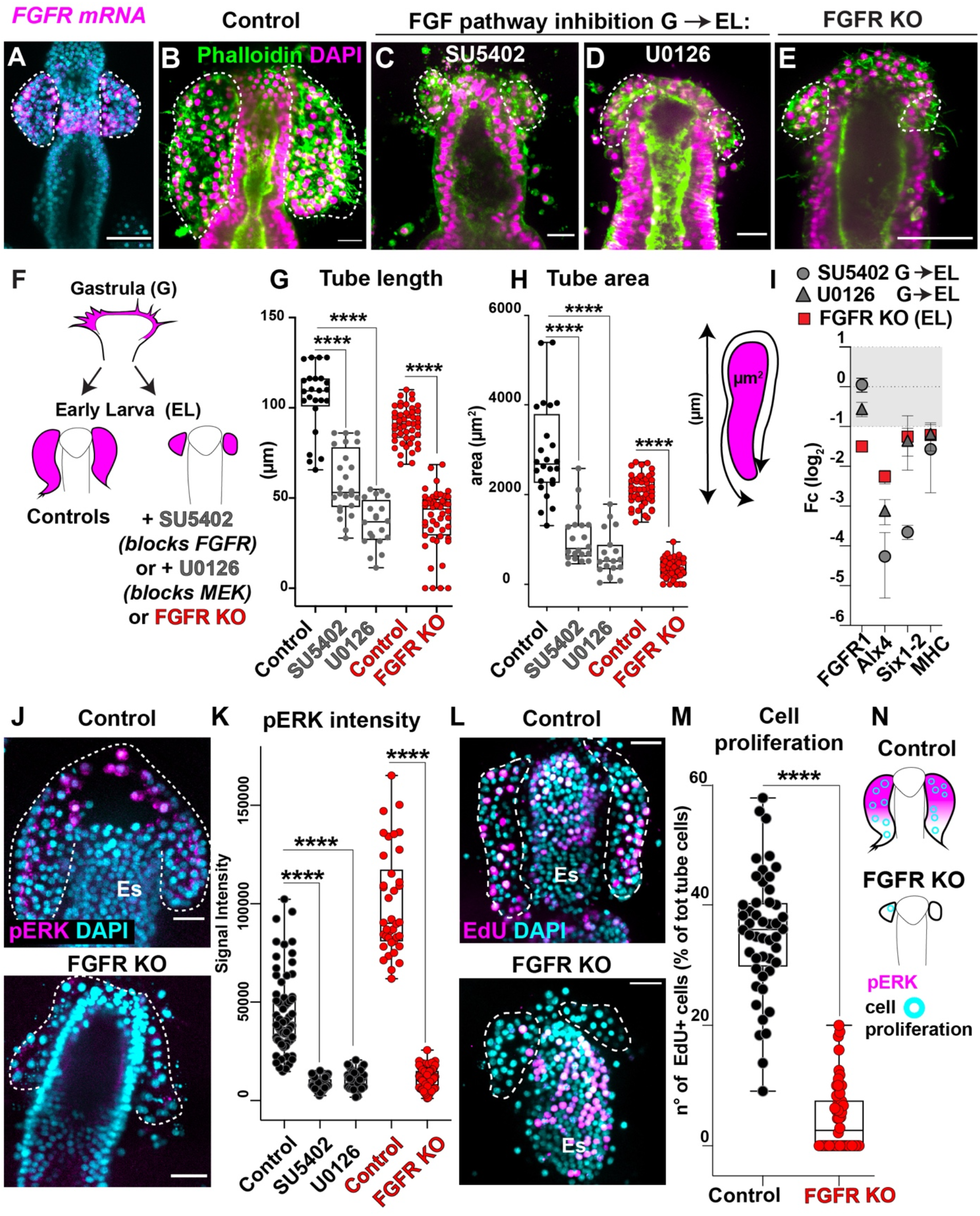
FGF signaling promotes tube outgrowth through pERK. **A)** FISH showing that FGFR is expressed in scattered tube cells. **B)** Control early larva (DMSO) and early larvae that were treated with **C)** the FGFR inhibitor SU5402 (final concentration 30μM) or **D)** the MEK inhibitor U0126 (final concentration 10μm) from the end of gastrulation to early larvae (GàEL) showing that both treatments lead to failure of tube outgrowth. **E)** Early larvae, in which the FGFR is knocked out by Cas9, also lack the tubes. **F)** Cartoon summarizing FGFR inhibition phenotypes. **G, H)** Tube length and area are significantly decreased in early larvae where the FGF pathway was blocked. Schematic showing how the length and the area of the tubes was measured. N= number of analyzed tubes (controls=26, SU5402= 26; U0126= 20, CAS9 control= 48; FGFR KO= 50). **I)** qPCR of significant genes in early larvae knocked out for FGFR or treated with SU5402 from gastrula stage show reduction of markers for tube cells. **J)** Phosphorylated ERK is active in control early larvae but absent in FGFR knock outs; **K)** signal intensity of pERK in the above experiments. The Y axis in the graph shows the raw integrated intensity for pERK normalized by the background signal for each image. **L)** EdU labeling (EdU pulse for 30 minutes) to mark proliferating cells that have recently synthesized DNA in control and FGFR KO early larvae. Es= esophagus. **M)** Quantification of EdU+ cells in controls and FGFR KO. Since the tubes of the FGFR KO larvae had fewer cells than controls, we assessed proliferation rate by counting the EdU+ cells normalized to the total number of tube cells (proliferation is represented as the % of Edu+ cells over the total number of cells in the tubes). Number of larvae n= 48 (controls), 50 (FGFR KO). **N)** Schematic showing that inhibition of FGF pathway at different levels prevents tube growth and pERK activation. For Student’s t-test ****p < 0.0001. White dotted lines outline the tubes. A, B, C, D, O, P scale bar is 50 μm. E, F, I, I’, j, K, L, M scale bar is 20 μm.

Knowing that FGF signaling promotes cell proliferation through pERK (Lovicu & McAvoy, 2001; Pagès et al., 1993), we tested whether FGFR influenced the cell proliferation we observed in the initial steps of tubulogenesis. We performed EdU pulse labeling to mark cells that have synthesized DNA in early FGFR knockout larvae and found a significant reduction of proliferating tube cells (Figure 6 L, M). This reduction in FGF-dependent cell proliferation was specific for the tube cells, since cells of the esophagus that also expressed FGFR (Figure 6 A) did not show pERK activity (Figure 6 J) and proliferation was not reduced there when FGFR is knocked out (Figure 6 L). We conclude that the FGF signaling pathway initiates tube outgrowth by promoting tube cell proliferation through pERK (Figure 6 N).

### Differential RNA-seq reveals genes induced by the FGF pathway during tubulogenesis

The FGF pathway is involved in multiple aspects of tubulogenesis, but the genes that link it to morphogenesis are still poorly understood (Bernascone et al., 2017). Blocking FGF signaling either genetically or pharmacologically precisely ablates the tube structures in sea star larvae (Figure 6 C, D, E). To define the landscape of genes activated by the FGF pathway, and potential genes that define cell types of the hydro-vascular organ, we generated tube-ablated larvae by SU5402 treatment soon after gastrulation. We then performed differential RNA-seq on these samples. In the analysis we selected candidates with an FDR SU5402/Controls <0.05 and fold change threshold of ±1.6 log_2_ SU5402/Controls, as used in previous studies (Czarkwiani et al., 2021; Wei et al., 2006; Weitzel et al., 2004). With this approach, our analysis identified 147 downregulated and 346 upregulated transcripts (Figure 7 A, Figure 7- Table 1, Figure 7- Figure Supplement 1).

**Figure 7.**
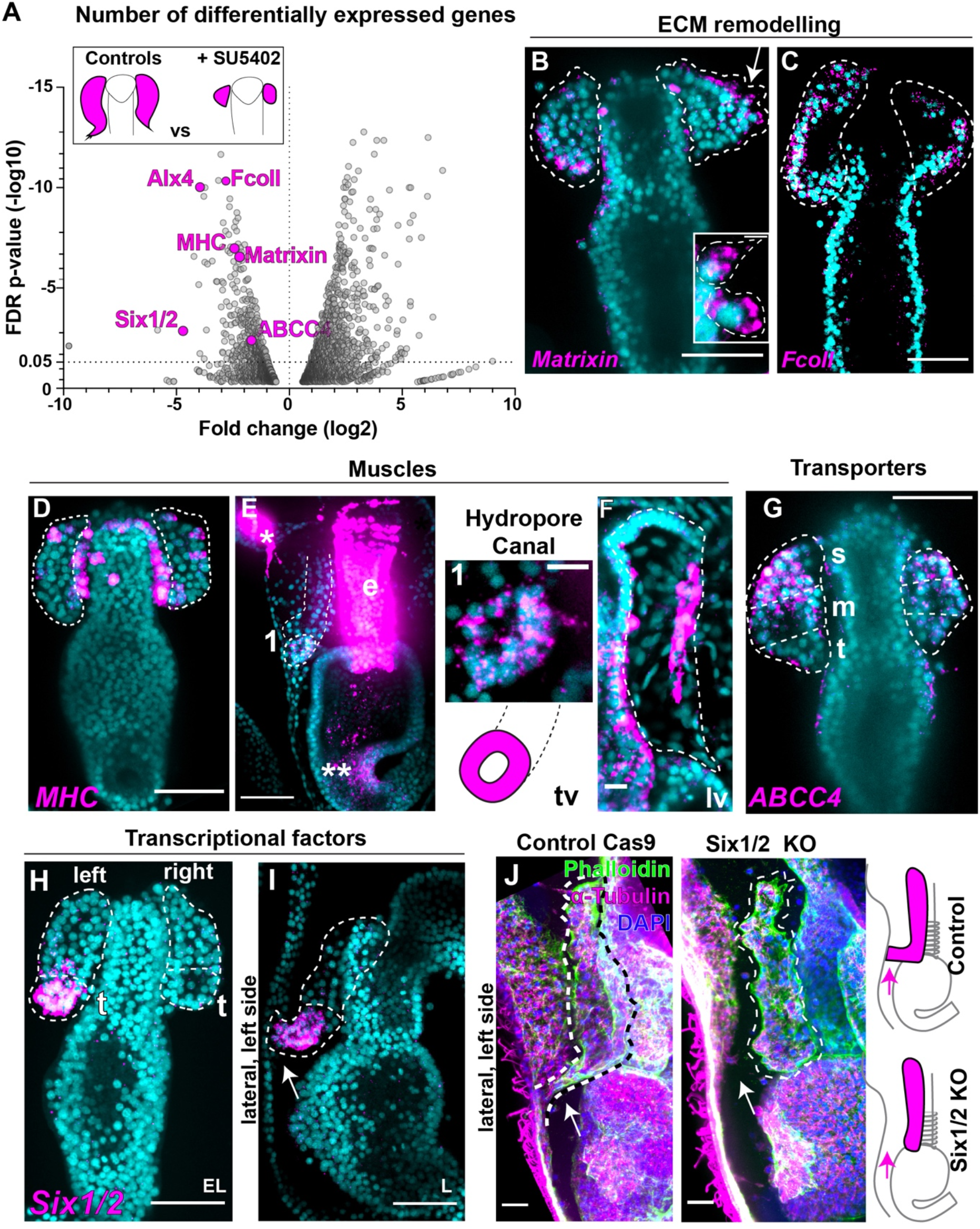
Differential RNA-seq reveals genes responsive to FGF signaling. **A)** Volcano plot of differentially expressed genes between controls and larvae where FGFR was blocked. **B, C)** FISH for genes (Matrixin, Fcoll) involved in extracellular matrix (ECM) remodeling. Arrow in B shows matrixin transcripts in cells extending protrusions. **D-F)** FISH showing myosin heavy chain (MHC) gene expression in the hydropore canal (1), the pyloric sphincter (** in E), the tube longitudinal muscle (F), the dorsal muscles (* in E) and the esophageal muscles (e in E). Tv= transverse view; Lv= lateral view. **G)** FISH for the transporter ABCC4 shows expression in tube stalk cells. **H, I)** FISH of Six1/2 showing gene expression at the tip of the left tube and later in the hydropore canal (arrow). **J)** Lack of hydropore canal in CRISPR Cas9 of Six1/2 compared to controls (arrow). All images are maximum projections of confocal z-stacks. s=stalk; m=medial; t=tip. B, C, D, E, G, H, I, J scale bar is 50 μm. E1, F scale bar is 10 μm.

As the SU5402 treatment inhibits tube formation, we focused on genes that were downregulated to test which candidates were uniquely expressed in the initial tubes. Most of the downregulated genes were involved in extracellular matrix homeostasis (Figure 7 -Figure Supplement 1). For example, we identified the extracellular protease matrixin as a target of FGFR, and its transcripts were enriched in lateral tube cells (Figure 7 A, Figure 7- Figure Supplement 2). Our RNA-seq study also identified the secreted the Kazal type serine protease inhibitor FcoII as a FGFR target, which we found to be enriched in the tubes (Figure 7 C, Figure 7- Figure Supplement 3). Another class of genes downregulated when FGFR was blocked were muscle-related genes, including troponin, myophilin, twist, myosin light and heavy chain (Figure 7_Figure Supplement 1). We found that the muscle marker myosin heavy chain (MHC) was initially expressed first by a few tube cells, but once the tubes elongated they formed two distinct populations of muscles (Figure 7 D, E, F). The first one was a muscle in the hydropore canal (figure E, insert 1). Here, MHC transcripts marked cells that did not have long fibers, similar to the intracellular contraction apparatus seen in the endodermal muscles of the sea star pyloric sphincter (** in Figure 7 E, 1) and in sea urchin larva pyloric and anal sphincters (Andrikou et al., 2013). The second muscle cells formed longitudinal fibers that appeared in L3 stage (Figure 7 F) and later extended along the whole length of the hydro-vascular organ (Figure 1 -Figure Supplement 1). Live image showed that the twitching of this longitudinal muscles allowed for the contraction of the hydro-vascular (Figure 7- Video 1).

Another group of genes downregulated in FGFR blocked larvae were transporters. The ATP-binding cassette sub-family C member 4 (MRP4 or ABCC4) is a multidrug resistance-associated protein whose expression in the mouse brain is dependent on FGFR1 (Jukkola et al., 2006). However, a role for ABCC4 in other morphogenetic mechanisms is unknown. We found that ABCC4 was a FGF target and was expressed in the initial tube, particularly in the stalk cells (Figure 7 G, Figure 7- Figure Supplement 4), suggesting that the link with the FGF pathway is conserved across species, even if the transporter is expressed in different tissues.

The transcription factors most affected by FGFR inhibition were Alx4 (Figure7- Figure supplement 5) and Six1/2. We found that in sea star larvae, Six1/2 was enriched in the tip of the left tube during tube elongation and later it marked the hydropore canal (Figure 7 H and I, arrow). Because we found that Six1/2 is expressed in a defined structure of the tube (the opening towards the outside environment), we focused on it for further investigation. Strikingly, we found that Six1/2 knockout larvae, while otherwise morphologically normal, did not form the hydropore canal, even 3 days after this structure normally forms in controls (Figure 7 J and knocked down in Figure 7- Figure supplement 6). Since the hydropore canal forms by tissue bending at a defined branch point, we propose that Six1/2 activation downstream of FGF signaling is critical for branching morphogenesis. In summary, blocking the FGFR pharmacologically we found a variety of candidate genes that are activated by the FGF pathway and that defined subpopulations of tube cells during tubulogenesis.

## Discussion

The tremendous diversity in strategies to form hollow tubes suggests that either these structures are not homologous, or that the toolkits to build such structures are not conserved. Here, we took advantage of the tubular hydro-vascular organ of the sea star larva to unravel the mechanisms of tubulogenesis in an intact, early branching deuterostome. We show that tube formation and elongation require that cells migrate and divide in defined zones. Cell fate depends on Notch signaling, while tube orientation requires Wnt signaling through the receptor Frizzled 1/2/7. In parallel, cell proliferation downstream of FGF signaling is critical for tube outgrowth and extension. Finally, we find that FGF reception induces the expression of genes potentially involved with hydro-vascular organ function, including muscles, ECM remodeling genes and transcription factors. These observations in the sea star, a representative of the sister group to all other deuterostomes including vertebrates, provide fundamental insight into the evolution of tubular organs.

### A transition in tubulogenesis strategy appeared in early branched deuterostomes

A fundamental feature of tubular organs is that the constituent epithelial cells are polarized. Tubes can form either from sheets of polarized cells or from nonpolarized cells that can acquire polarization during morphogenesis to form a lumen *de novo* (Hogan & Kolodziej, 2002). We found that the hydro-vascular organ develops from pre-polarized precursor cells, therefore positioning this organ among the first category. Figure 8 A summarizes the main steps of the sea star tubular organ development. Mesodermal cells from the tip of the growing gut during gastrulation evaginate to form two initial tubes formed by a simple monolayer epithelium that keeps its polarity during growth. Cells are ciliated and rich in actin at the apical side facing the lumen, while the basal side faces the basal lamina (Figure 1 D-G). Similarly, organs like the Drosophila salivary glands and vertebrate lungs also form from polarized precursors that migrate to accomplish tube morphogenesis (Chung & Andrew, 2008; Hogan & Kolodziej, 2002; Lubarsky & Krasnow, 2003). However, the mechanisms of cell dynamics in tubulogenesis differ between *Drosophila* and mammalian tubular organs. For instance, *Drosophila* salivary gland and trachea precursor cells complete proliferation prior to beginning tubular morphogenesis (Kerman et al., 2006; Kerman et al., 2008; Samakovlis et al., 1996; Skaer, 1989), while in vertebrate organs like lungs and kidney cell proliferation and migration are coupled (Hogan & Kolodziej, 2002; Mollard & Dziadek, 1998; Vasilyev et al., 2012; Weaver et al., 2000). Is this a generalizable difference between vertebrates and invertebrates? Or alternatively, does this difference instead reflect the consequence of distinct life histories between individual species?

**Figure 8.**
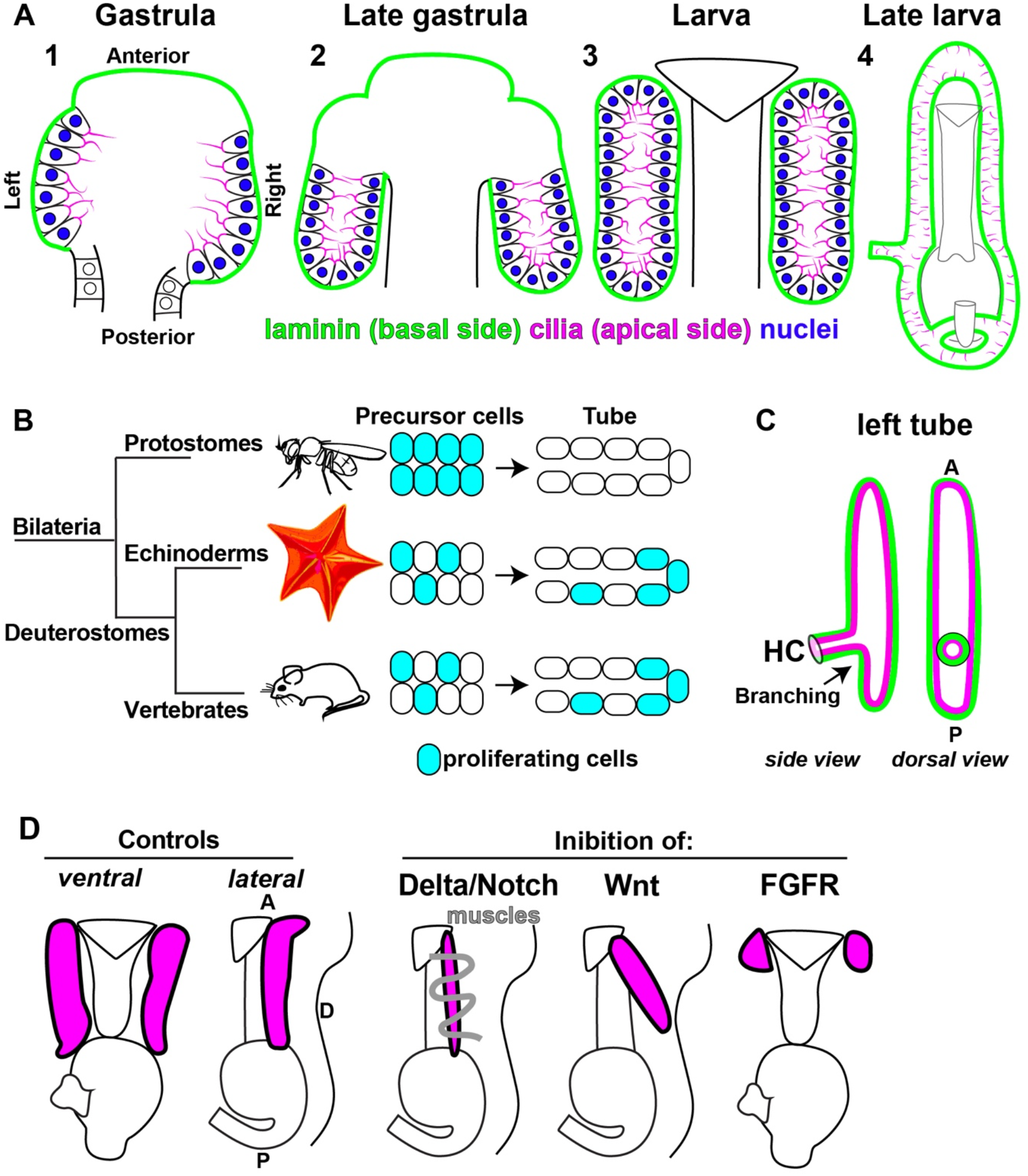
Summary of the intrinsic and extrinsic factors that guide tubulogenesis of the hydro-vascular organ. **A)** Stages of sea star hydro-vascular organ development. The precursor cells of the hydro-vascular organ come from mesodermal progenitors at the tip of the growing gut (1). In the first stage of tubulogenesis, pre-polarized cells form a lumen and grow towards the posterior side of the embryo to form two tubes (2). During embryogenesis, left and right tubes detach from the esophagus (3) and finally form a close system where the left and right tube connect anteriorly and posteriorly, the hydro-vascular organ. **B)** Phylogenetic tree showing the relationship of the *in vivo* systems for the study of tubulogenesis. In protostomes tubular organs form by a first step of cell proliferation followed by morphogenesis. In echinoderms and in vertebrates (both deuterostomes) cell proliferation is coupled with cell migration during morphogenesis. **C)** cartoon showing the branch formed by the left tube, the HC (hydropore canal) that opens on the outside environment. **D)** Cartoon summarizing the role of the signaling pathways on tube formation. A, anterior; P, posterior; D, dorsal.

Here we show that as in vertebrates, cell migration and cell proliferation are coupled during tubulogenesis in the sea star, an early-branched deuterostome (Figure 8 B). To establish the initial tube, cells undergo a two-phase migration from the tip of the growing gut to finally run parallel to the gut itself (Figure 2 A-H). Similarly to tubulogenesis in vertebrates, subsets of cells actively divide during these migrations in the sea star. Localized cell proliferation is an important driver of shape changes, as in mammalian tooth, mammary gland, and embryonic lung morphogenesis (Daniel et al., 1984; Goldin et al., 1984; Jernvall & Thesleff, 2000). In the sea star tubes, we find that proliferation is downstream of FGF signaling, and it is concentrated in two main proliferation zones: the middle region of both tubes and the tip of the left tube. We propose that the extensive proliferation in the middle of the tube drives A-P elongation. Instead, proliferation at the tip of the left tube may drive formation of the critical branch point that forms the hydropore canal (Figure 8 C). Indeed, by blocking cell cycle progression by Cdk4/6 inhibition, or FGF perturbations at different time points, tube initiation or elongation was disrupted. FGF signaling regulates proliferation in the mouse Wolffian duct (Okazawa et al., 2015), in the kidney metanephric mesenchyme (Poladia et al., 2006), and in the mammary gland (Pond et al., 2013). Similarly, the FGF signaling pathway is required for the formation of homologous tubular pouches in the sea urchin, a related echinoderm, suggesting that these mechanisms are conserved within this clade (Andrikou et al., 2015). These striking similarities with vertebrate morphogenesis suggest that the coupling of proliferation with cell migration may be an ancestral deuterostome innovation for the formation of tubular organs.

### Distinct extrinsic signals regulate elongation, cell fate and orientation of tubes

The shape of epithelial tubes is guided by signaling pathways from the embryonic environment. However, because of the great morphological complexity of tubular organs, our knowledge on the contribution of the extrinsic cues to distinct steps of tubular organ establishment is still limited (Bernascone et al., 2017). The sea star hydro-vascular organ offers a simplified system in which we defined how the Delta/Notch, Wnt and FGF pathways each uniquely contribute to distinct aspects of tubulogenesis (summarized in Figure 8 D).

Tube cells share a common origin with muscles, as both tissues derive from the mesoderm (Andrikou et al., 2015; Barone et al., 2022; Cary et al., 2020; Hinman & Davidson, 2007; Sherwood & McClay, 1999). Here we report that when we block Delta/Notch signaling, an important regulator of mesodermal specification (Materna et al., 2013), excess muscle fibers enwrap the tubes causing constriction (Figure 4 E-G). There are two possible hypotheses on the origins of these fibers. First, Delta/Notch signaling acts on cells from the adjacent esophageal muscles to induce migration to the tubes and differentiation once there. Alternatively, our data support that Delta/Notch signaling might inhibit epithelia tube cell transfating into a muscle-like phenotype. Notch inhibits skeletal muscle differentiation in many contexts, including during vascular system differentiation (Gerrard et al., 2021; Gioftsidi et al., 2022; D. Morrow et al., 2008; Pardo-Saganta et al., 2015; Shawber et al., 1996). From our data, these ectopic muscle fibers appear to originate from tube epithelial cells (Figure 4 F, G). Moreover, we observed a few cells from the tubes undergoing EMT to become mesenchymal (Figure 2 B and Figure 2- Video 1), suggesting that some cells retained the ability to acquire a distinct mesodermal fate. This phenotypic change, that eventually leads to loss of epithelial function, has been observed in lungs and kidneys to be the cause of renal and lung fibrosis (Simon & Hertig, 2015; Willis et al., 2006), but it is unclear what pathways guide it. Therefore, our data in the sea star suggests that the Delta/Notch signaling might be responsible for such phenotypic change in complex organs.

Although correct orientation is critical for tube function, very little is known about how tubular organs orient within the growing embryo (Bernascone et al., 2017). Taking advantage of the simple morphology of the sea star tubes, we found that the Wnt signaling through the Frizzled 1/2/7 receptor regulates tube orientation (Figure 5A-G and 8D). Importantly, however, gene expression for tube markers was not affected by lack of Wnt signaling (Figure 5-Figure supplement 1). We therefore hypothesize that Wnt signaling may be acting through the planar cell polarity pathway rather than the canonical pathway to guide shape changes and tube directionality. In support of this premise, Wnt signaling has been found to coordinate cell polarity in groups of cells, for instance by controlled cell division orientation in the mouse stratified epithelium (Box et al., 2019; Yamamoto et al., 2011). Moreover, the Frizzled 2 receptor controls branch formation and shape of the epithelial tube of the highly branched mouse lungs (Kadzik et al., 2014). These findings all highlight the conserved importance of Wnt signaling in defining the shape and orientation of complex organs across evolution. Importantly, our study adds that Wnt signaling specifically guides the 3D orientation of the growing tube in embryogenesis.

### Genes induced by the FGF pathway reveal a breadth of mechanistic insight

The downstream targets of signaling pathways in tubulogenesis are poorly understood (Bernascone et al., 2017) and the finding that FGF perturbation selectively ablated the tubes allowed us to use RNAseq to discover putative transcriptional targets of the FGF pathway in the developing hydro-vascular organ. The most abundant candidates we found were metalloproteases and tissue inhibitors of metalloproteases (expression examples for matrixin and FcoII, Figure 7 B, C). Dynamic remodeling of the extracellular matrix components (ECM) at the basal and apical compartments of tubes is a fundamental factor in tube shape and growth (Loganathan et al., 2020). The basal lamina controls the size of the Drosophila tracheal tubes (Klußmann-Fricke et al., 2022), and remodeling of the ECM through metalloproteases is essential for the elongation of the sea urchin larva skeleton, a syncytial tube (Morgulis et al., 2021). Since matrixin and FcoII proteins have a signal peptide it is possible the metalloprotease and its inhibitors are secreted to contribute to ECM remodeling around the growing tube. The balance of these factors is regulated by the FGF pathway. In further support of this model, tube cell precursors are enriched in genes related to the extracellular matrix and cell/cell adhesion (Meyer et al., 2022).

Another group of FGF downstream candidates were muscle genes, and we focused on the expression of the terminal differentiation gene MHC since it is highly abundant. We discovered two new populations of muscles not previously described in the sea star larva: muscles in the hydropore canal and longitudinal muscle fibers along the tubes (Figure 7 D-E). The longitudinal muscle allows for the tubular organ to contract (Figure 7_Video 1), while the movements of cilia inside the lumen helps moving fluids or particles in the tubes (Figure 1 -Video 3). Studies in other echinoderms showed that larvae have smooth muscles (Burke & Alvarez, 1988), suggesting this is also the case of the longitudinal muscles around the hydro-vascular system. Our findings show that the hydro-vascular organ also includes muscles that surround this organ in a way that resembles the vasculature or the intestine, specialized tubes that can autonomously contract to move their contents forward.

### A conserved regulatory network for branching morphogenesis

In our RNAseq analysis of FGF targets, we identified the transcription factor Six1/2, which is selectively expressed in the hydropore canal (Figure 7 H-J). By Cas9 knockout, we found it to be essential and selective for forming the hydropore canal branch point. In earlier stages, Six1/2 was found to be expressed in the mesodermal tube precursor cells in both sea stars and sea urchins (Andrikou et al., 2015; Cary et al., 2020; Koop et al., 2017; Martik & McClay, 2015; Yankura et al., 2010). In an example of potentially homologous function, in the mouse, Six1 plays a critical role in a subpopulation of collecting tubule epithelial cells in the developing kidney (Xu et al., 2003). Six1/2 is also expressed by distal epithelial tips of branching tubules in embryonic lungs where it coordinates FGF signaling (El-Hashash et al., 2011). In addition, FGF signaling regulates branching by activating Six1/2 in the lachrymal gland (Garg et al., 2018). These striking parallels between formation of the branch point in the sea star hydro-vascular organ and different examples in mice suggest that Six1/2 activation by FGF may be a conserved node in the regulation of branching morphogenesis. An advantage of the sea star system is that while mammalian organs have multiple branches, the sea star tubes offer a simple system with a single defined branched point observable in vivo (Figure 8 C). This system will enable future studies to unravel what aspects of branching are specifically regulated by Six1/2 and its interactions with FGF signaling.

### The sea star hydro-vascular organ offers insight into organ evolution

Our results support the idea that the basal toolkit to make vertebrate tubular organs was already established at the root of the deuterostome clade, the group that includes vertebrates. We revealed that, as in vertebrate organs, tubulogenesis is driven by cell migration coupled with proliferation in the sea star hydro-vascular organ. Like in vertebrate organs, proliferation happens in growth zones, especially at the tube tips, and it is regulated by FGF signaling. Six1/2 is a conserved target of FGF signaling that is critical for establishing branch points. Given the simplified morphology of the sea star tubes compared to highly branched vertebrate organs, and the technical advantages that the system offers for *in vivo* studies, we were able to discover new roles of signaling pathways in tubulogenesis. We propose that the development of the sea star hydro-vascular organ provides important insight into the fundamental evolutionary steps that preceded the appearance of the more complex vertebrate organs.

## Materials and Methods

### Embryo and larva cultures

*Patiria miniata* adult animals were obtained from Pete Halmay and Marinus Scientific and kept at 15°C in a sea water aquaria. Gonads were surgically obtained by small cuts on the oral side. Embryos and larvae were cultured at 16°C. Oocytes were microinjected, then treated with 1-methyladenine (Acros Organics) at a final concentration of 10 μM to induce meiotic resumption.

### Plasmid constructs and RNA in vitro transcription

The H2B-GFP (pZS317) construct was amplified from first strand cDNA reverse transcribed from total ovary mRNA and then cloned into pCS2+8 as C-terminal GFP fusions (Gökirmak et al., 2012). The plasmid was linearized with NotI to obtain a linear template DNA. mRNA was transcribed in vitro with the mMessage mMachine SP6 followed by the polyadenylation kit (Life Technologies), then purified using lithium chloride. In vitro transcribed mRNA was injected at a final concentration of 500 ng/ul.

### Perturbation experiments with MASO injection and CRISPR Cas9

Prophase arrested oocytes were injected with approximately 20 picoliters of H2B mRNA, Six1/2 morpholino antisense oligonucleotide (MASO) or Cas9+gRNAs solutions in nuclease free water. A MASO (GeneTools, Philomath, OR) complementary to *Pm-Six1/2* transcript 5’ UTR was used at a final concentration of 800 μM to block protein translation (5’-CGTGAAGCCAAACGACGGCAACATG-3’; as used in Cary et al., 2020). A scrambled MASO sequence was used as control. For each knock out experiment, three Cas9 guide RNAs (gRNAs; 150 ng/μl of each gRNA) were used to target genomic DNA as previously described (Perillo et al., 2022). gRNAs (Synthego) were mixed with 750 ng/μl of Cas9 mRNA; control larvae were injected with Cas9 mRNA only. Guide RNAs sequences are: Pm-Delta: 519_AACGGAGGCACCTGCGAGAA; 198_CGGCCCAAGAACGACAGCTT; 248_GAAAGTGTGCTTGGATGGCT. Pm-Six1/2: 44_CTGCGAAGTTTTGCAGCAGT;86_CCGGCAGCGACCAGAGAAAA; 274_AGGCTGAGAAACTCCGGGGC.

Pm-FGFR1: 185_GGCGTCGTGGTAGAAGACAG; 136_GGAGACGGGGGATGACAGAG; 31_ GGTCACAGAACTGAAGAAAG.

Pm_Fz1/2/7: 27_ACGGTGTGGGCCGTGAAATC; 81_TCTGCGGTCGAATGTCAAAT; 200_CAGACAGGAGGAAGCCGGCT.

Genomic DNA of individual larvae was sequenced to test for CRISPR Cas9 induced mutations as previously described (Oulhen et al., 2022). Genomic sequences used to find *Patiria miniata* genes and to design gRNAs were analyzed with the resources at echinobase.org (Arshinoff et al., 2021).

### Live imaging, single nucleus tracking and statistical analyses

For time-lapse recordings, larvae were embedded in 1% low melting point agarose (Sigma), covered with sea water, mounted on a MatTek 35mm glass bottom dish, and imaged at room temperature (22°C). For tube elongation movies, larvae of two independent experiments expressing H2B-GFP were imaged for 2 days capturing z-stacks with 3 μm step size every 5 minutes. For each experiment, both the left and the right tubes were imaged, and nuclei were tracked. Movies and images of fix samples were taken on a Nikon Yokogawa W1 spinning disk microscope with 40x silicon immersion oil objective. Images were processed and analyzed in Fiji (ImageJ) by using the manual tracking plugin (Fabrice Cordelières, Institut Curie, Orsay). To track the displacement of individual nuclei, we first normalized the movies by defining the starting point of tube elongation (t=0) as the tubes were 50±5μm long, and the final point of tracking (t=40-45h) as the tubes were 80±10μm long. Only nuclei of the tube epithelial cells that were in focus and moving along the x and y axes were tracked. Bright field movies were taken with a Zeiss Axio Vert.A1 in continuous mode. DIC video of cilia moving in the hydro-vascular organ were taken on an Olympus FV3000 confocal microscope (Brown University Leduc Facility). Student’s t-test was performed to assess statistical significance, values of p<0.001 are represented as ****. Graphs and statistical analyses were performed using Prism (GraphPad Software). Figures were made with Adobe Illustrator.

### EdU incorporation and pharmacological perturbations

5-ethynyl-2-deoxyuridine (EdU) pulse labeling was performed with the Click-IT EdU imaging kit (Life Technologies; Cat#:C10340) as described in (Perillo et al., 2022). A final concentration of 10 mM EdU in sea water for 30 min was used. 25 to 50 tubes were analyzed for each stage.

The Cdk4/6 inhibitor Abemaciclib (LY2835219) (Palumbo et al., 2019) was added to larval cultures at 16 °C at a final concentration of 10μM. The γ-secretase inhibitor DAPT (N-[N-(3,5-difluorophenacetyl)-L-alanyl]-S-phenylglycine t-butyl ester, FisherScientific) (Materna et al., 2013) was added to a final concentration of 10 μM. To perturb the Wnt pathway, the small molecule porcupine inhibitor ETC-159 (Katoh, 2017) was used at a final concentration of 5μM. The FGFR inhibitor SU5402 was used at 40μM final concentration and the MEK inhibitor U0126 at 10μM. For all pharmacological experiments, drugs were dissolved in DMSO and the optimal drug concentration was chosen after testing concentration ranges from 0.1 to 100μM. All drugs were added at the end of gastrulation, prior to the development of the hydro-vascular organ tubes. A corresponding volume of DMSO was added to sea water in controls. Experiments were performed in three independent biological replicates.

### Fluorescent *in situ* hybridization (FISH) and immunofluorescence

Larvae were fixed in 4% paraformaldehyde overnight at 4°C and fluorescence in situ hybridization (FISH) was performed as described in (Perillo et al., 2021). For immunofluorescence experiments, larvae were fixed in 2% PFA in PBS for 20 minutes at RT followed by 10 minutes in 100% methanol (this step was skipped only for experiments where phalloidin was added). Larvae were then washed in PBST (0.1% Tween20) and incubated overnight at 4°C with 1 mg/ml BSA and 4% sheep serum in PBST and one or more of the following: Alexa-fluor 488 Phalloidin 1:300 (Molecular Probes), anti-beta tubulin antibody 1:100 (E7, Hybridoma bank), anti-laminin antibody 1:300 (AB11575, Abcam), monoclonal Anti-MAP Kinase, Activated (Diphosphorylated ERK-1&2) pERK 1:100 (M8159, Millipore Sigma). Samples were then washed three times with PBST and incubated with the corresponding AlexaFluo secondary antibodies (Invitrogen) at a final concentration of 1:2000 for 2h RT. Samples were imaged on a Nikon Yokogawa W1 spinning disk microscope and processed in Fiji.

### RNA extraction and qPCR

RNA from 100 larvae was isolated with the RNeasy Micro kit (Qiagen, Cat#:74004). cDNA synthesis was performed using the Maxima kit (Life Technologies, Cat#:K1641). qPCR was performed with Maxima SYBR master mix (Life Technologies, Cat#:FERK0222) using ABI7900 and QuantStudio3 (ThermoFisher) real-time PCR systems. Transcripts were normalized to ubiquitin. Three biological replicates and three technical replicates were performed. A threshold of twofold difference was chosen as a biologically meaningful change. Points on the graphs show the mean of the three technical replicates and error bars represent biological replicates.

### Differential RNAseq

Total RNA was isolated from 3 day-old larvae (LG) stored in Trizol (Ambion) in three biological replicates. Samples were sent for RNA extraction, library preparation, Illumina HiSeq 2×150 bp sequencing, quality control and RNA-Seq data analysis to Genewiz (www.genewiz.com).

## Supplementary material legends

**Figure 1-Table 1.** Staging of hydro-vascular organ development.

**Figure 1- Video 1. Live imaging of tubulogenesis in the sea star embryo over the first 15 hours of tubular morphogenesis.** Time lapse of a live embryo developing from gastrula to early larva. Asterisk indicates the lumen of the newly formed tubes. Time is displayed in hours:minutes:seconds; video acquired with bright field microscopy.

**Figure 1- Video 2. 3D movie of a sea star late larva.** 3D reconstruction of a late larva from the ectoderm towards the internal organs. The main structures are annotated. Dashed line arrow indicates where the tubes meet at the anterior side to create a close system. A= anterior, P= posterior. Image taken with a 20x objective.

**Figure 1 -Video 3. DIC live imaging of cilia beating inside the larva left tube.** Time in min:sec.

**Figure 2 -Video 1**. Movie 1; cell tracking of nuclei of stalk cells. For all videos time is represented as hours:minutes and nuclei are marked by the probe H2B-GFP. Movies are maximum projections.

**Figure 2 -Video 2.** Movie 1; cell tracking of nuclei of left tube cells

**Figure 2 -Video 3.** Movie 1; cell tracking of nuclei of right tube cells

**Figure 2 -Video 4.** Movie 2; cell tracking of nuclei of tube cells (stalk, left and right).

**Figure 7- Video 1. Contraction of the tubes in a late larva.** Bright field movie of a live sea star late larva (20x objective) showing that the tube organ is vital and contracts thanks to the tube longitudinal muscles. Dashed line arrows indicate where muscle twitch happens. Particles in green are algae in the stomach. Time is indicated in seconds.

**Figure 1-Table 1.** Differential RNA-seq expression of larvae treated with SU5402 versus Controls; GOTerm ID for up and downregulated genes.

## Acknowledgments

The authors are grateful for the support for this work from the Eunice Kennedy Shriver National Institute of Child Health and Human Development (K99HD099315) to S.Z.S and the National Institutes of Health 1R35GM140897 to GMW. The authors thank Mary Jane Tsang and Iain Cheeseman for sharing reagents and PRS for technical assistance. We thank Synthego for providing free samples of FGFR gRNAs.

---

**Figure 1- Figure supplement 1.**
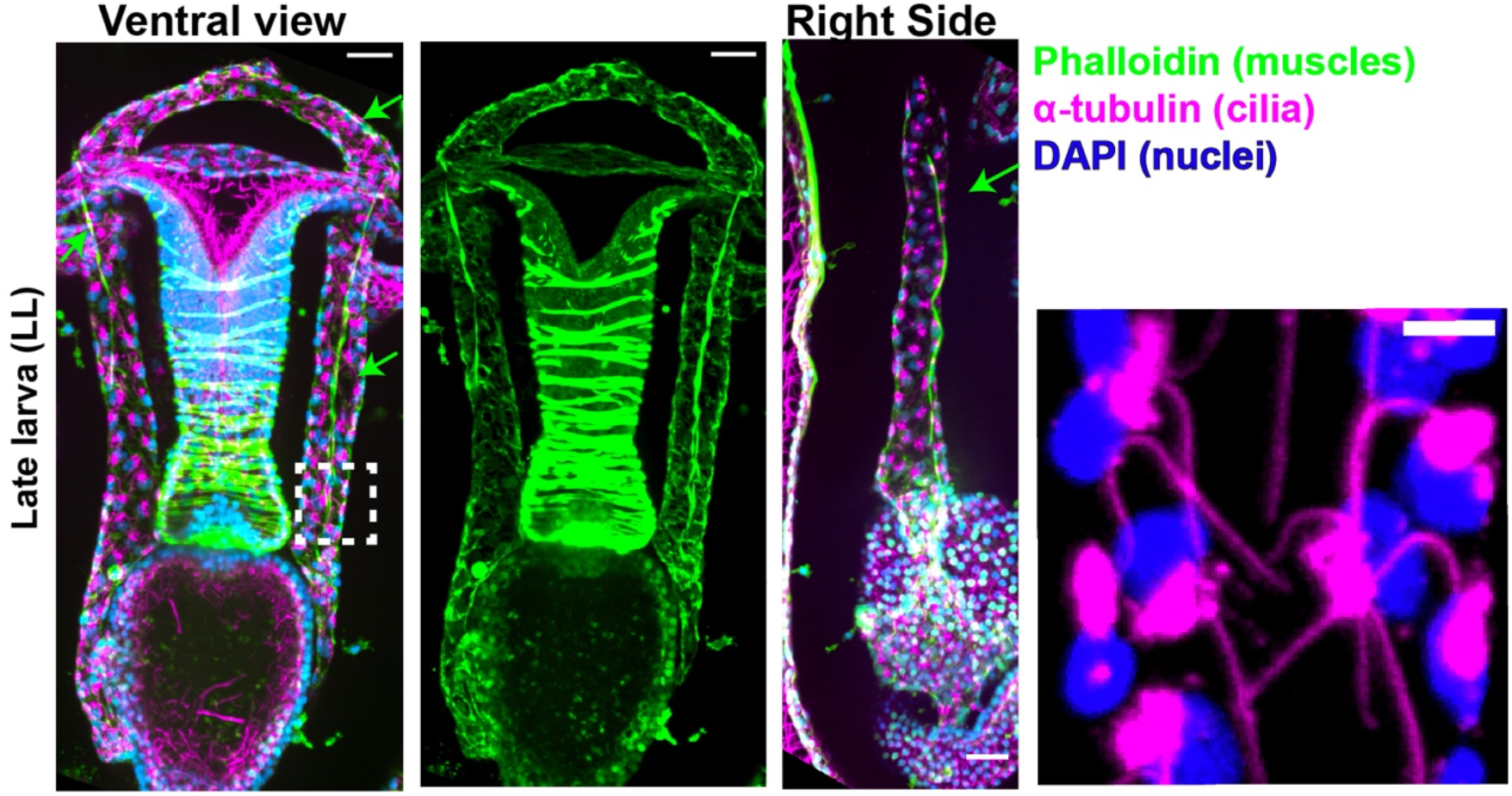
Alpha tubulin staining of a late larva showing that the epithelial cells of the tubes are ciliated (insert). Phalloidin (actin) label reveals extensive longitudinal muscles (arrows) that extend the most anterior and posterior tubular sites. Scale bar 20 μm.

**Figure 2 -Figure supplement 1.**
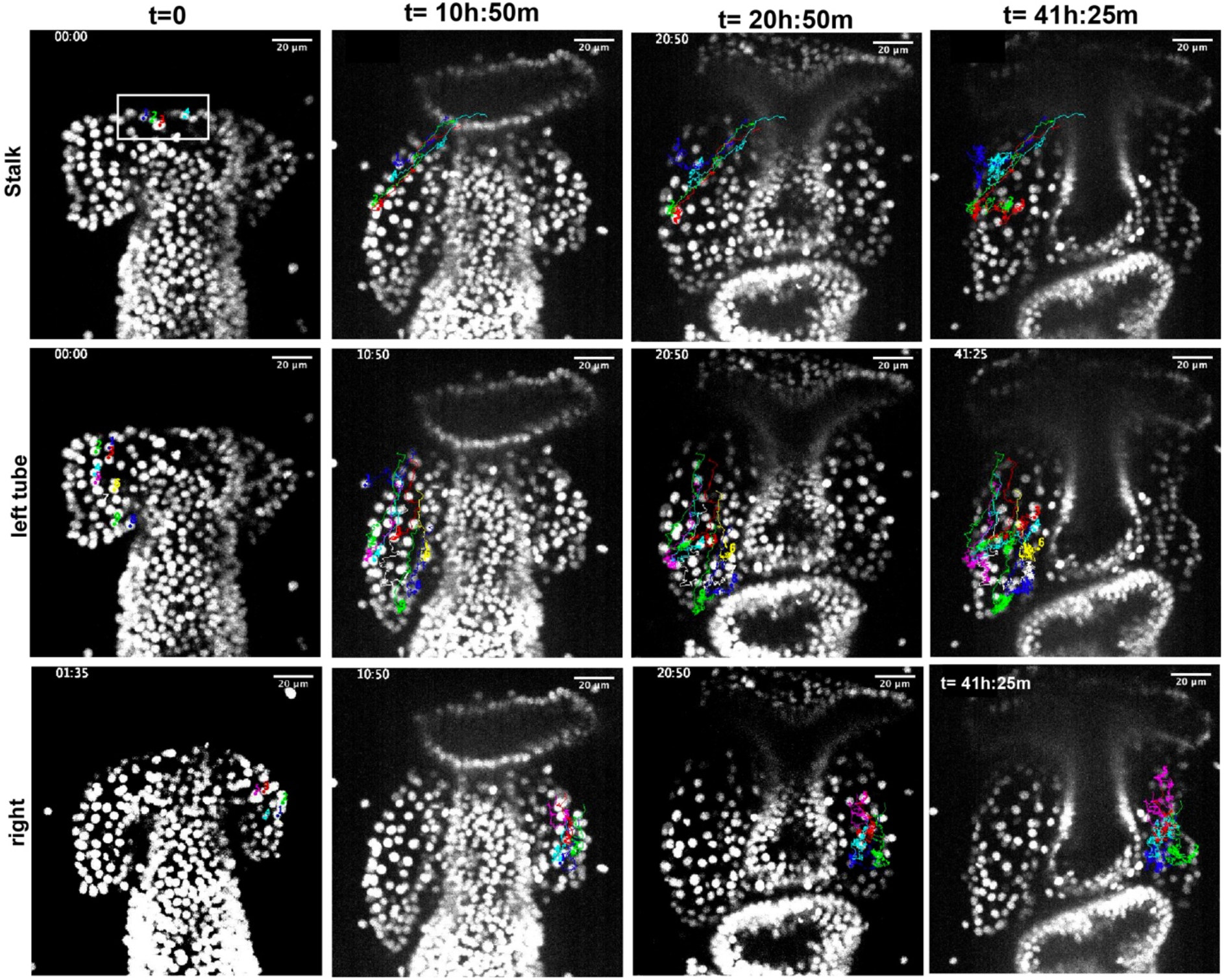
Time-lapse images of cell tracking for video 1 (still images from Figure 2- Video 4, 5 and 6). Tracking of cells of the left tube is shown in main Figure 2 A-D.

**Figure 3 -Figure supplement 1.**
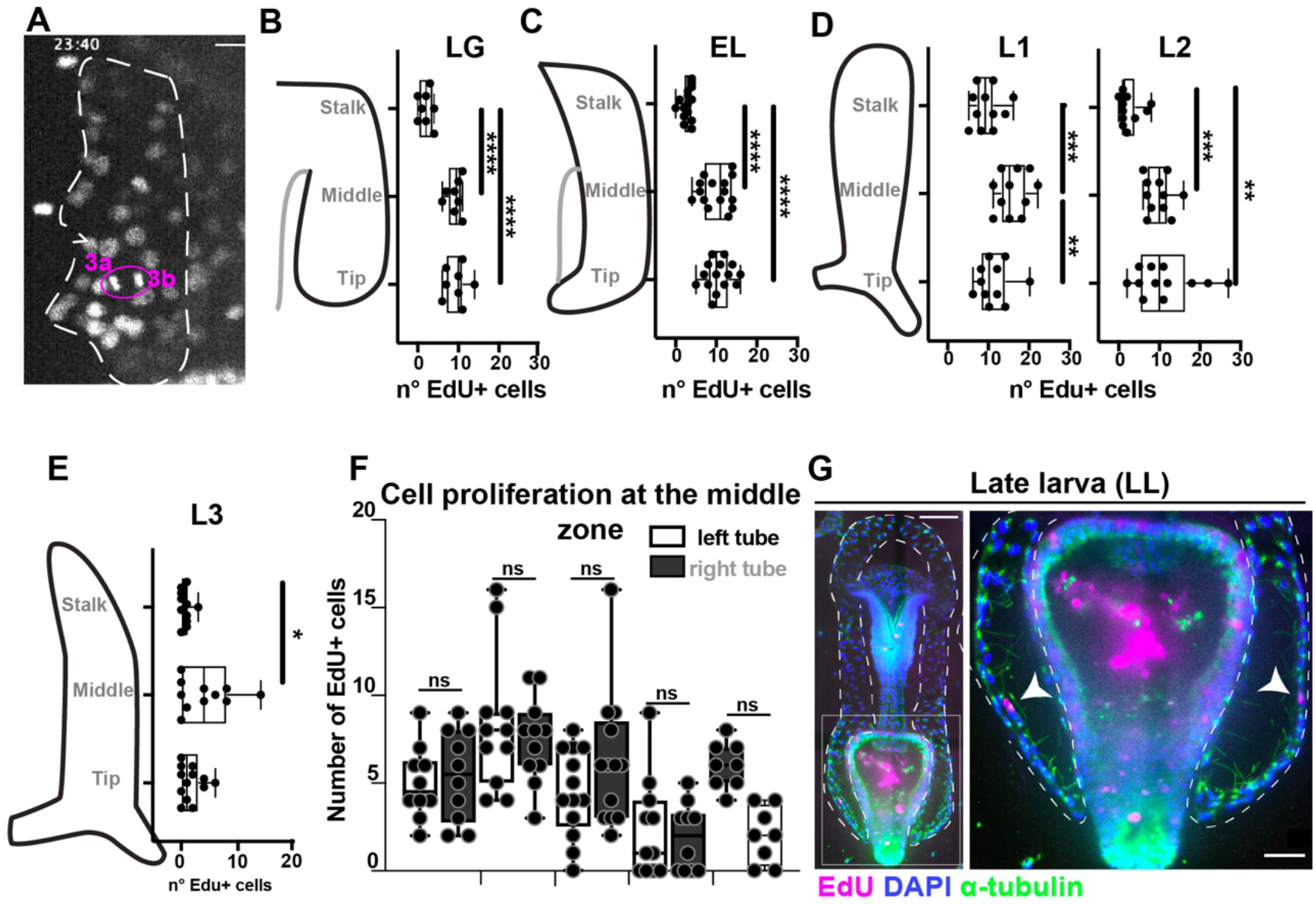
**A)** Single image frame from movie 1 showing a mitosis event. **B-E)** EdU+ cells across 3 tube zones: stalk, middle and tip. Graphs represent the data shown in the heat map of Figure 3 E. **F)** Cell proliferation of the middle zone is the same in left and right tubes. **G)** EdU incorporation in late larva showing lack of cell division in the tubes, except the left and right somatocoels (the posterior most-end of the tubes).

**Figure 4 -Figure supplement 1.**
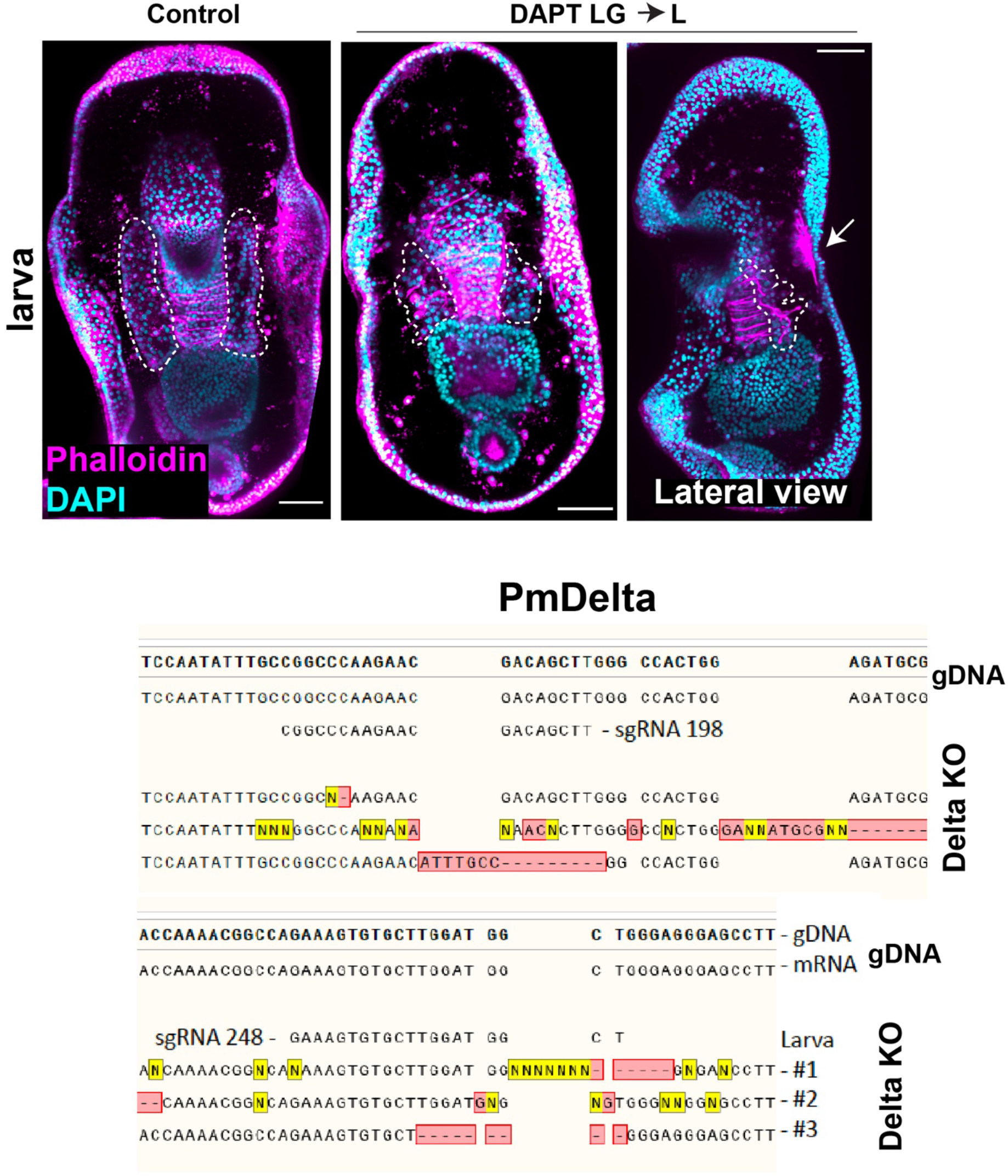
Treatments with the Delta-Notch inhibitor DAPT added at the end of gastrulation shows that tubes are shorter but overall larval morphology is intact. Scale bar = 50 μm. Sequencing results for Delta KO showing mutation in the genomic DNA.

**Figure 5- Figure supplement 1.**
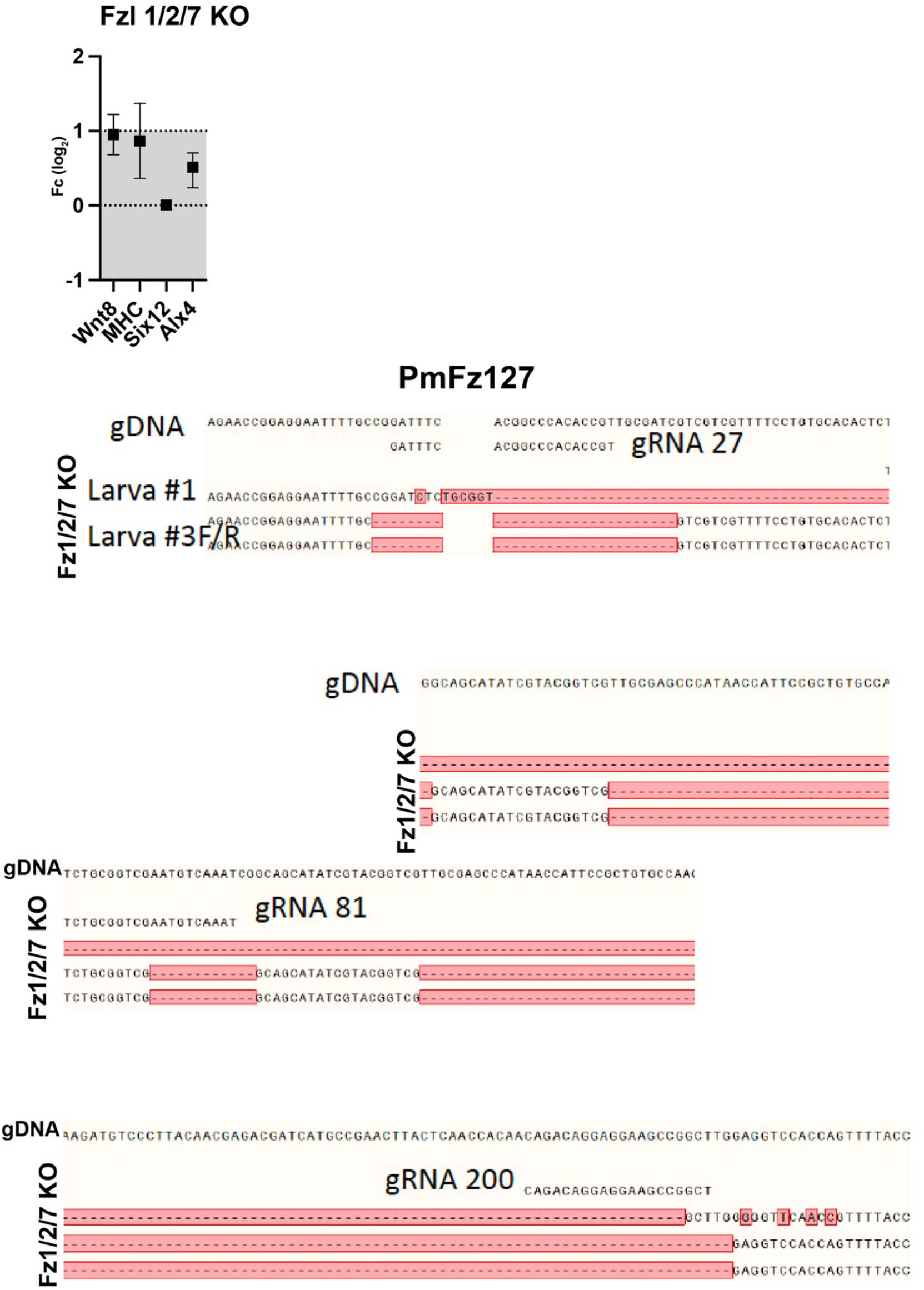
qPCR showing that, as a control for the ETC19 treatment, Wnt8 expression was decreased. Expression of tube genes and muscle markers was unchanged. Sequencing results for Fz1/2/7 KO showing mutation in the genomic DNA.

**Figure 6- Figure supplement 1.**
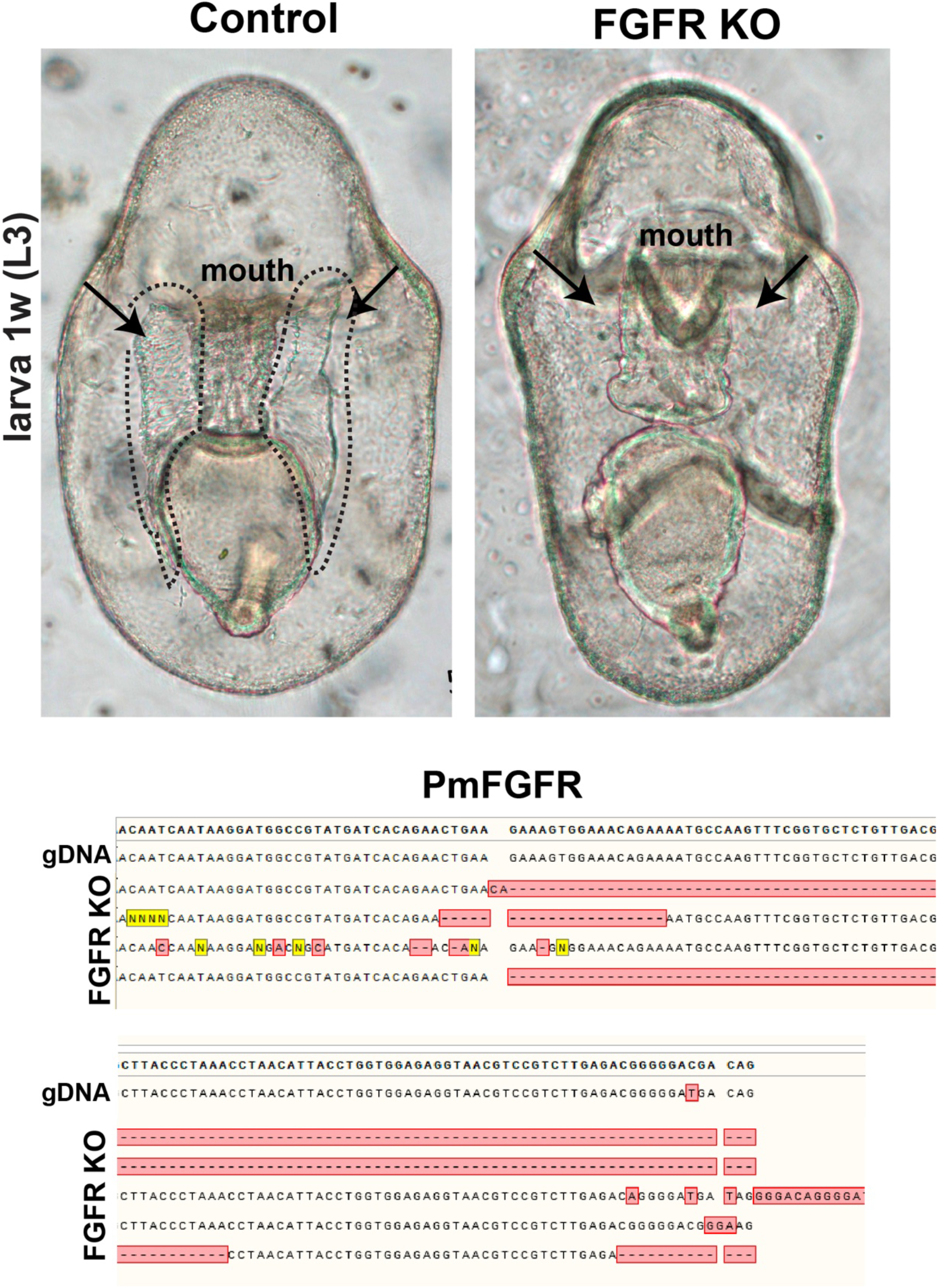
Controls larvae and larvae knocked out for FGFR show absence of tubes. Sequencing results for FGFR KO showing mutation in the genomic DNA.

**Figure 6- Figure supplement 2.**
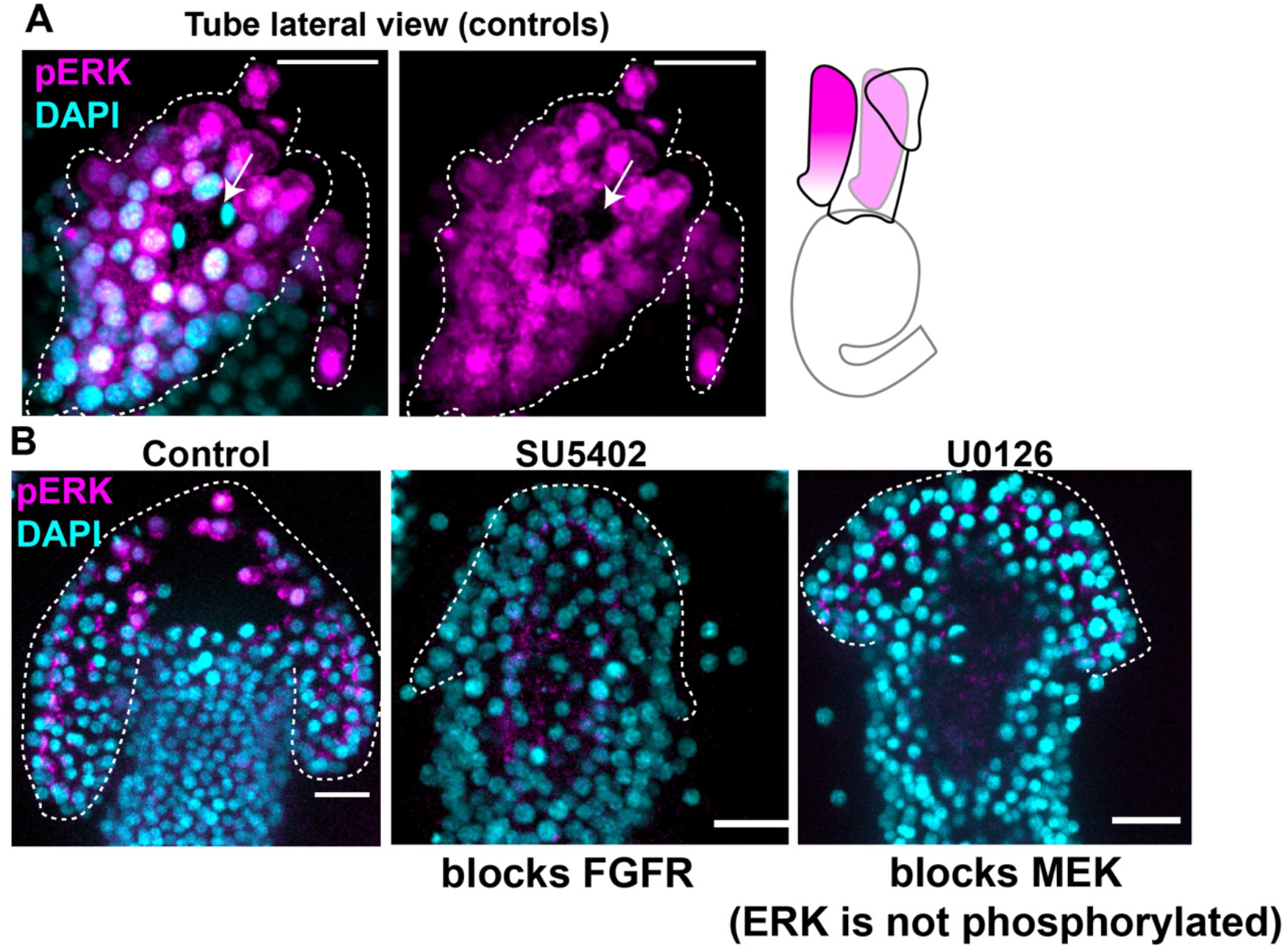
**A)** Immunofluorescence using the anti-phosphorylated ERK antibody showing that active ERK is localized in the nuclei of most tube cells (side view). Arrows showing that dividing cells do not have pERK staining. **B)** Immunolabeling of phosphorylated ERK showing that pERK is active in control early larvae but absent when the FGFR tyrosine kinase activity is blocked with the drug SU5402 and when MEK (the kinase that activates ERK) is blocked with the drug U0126.

**Figure 7 -Figure supplement 1.**
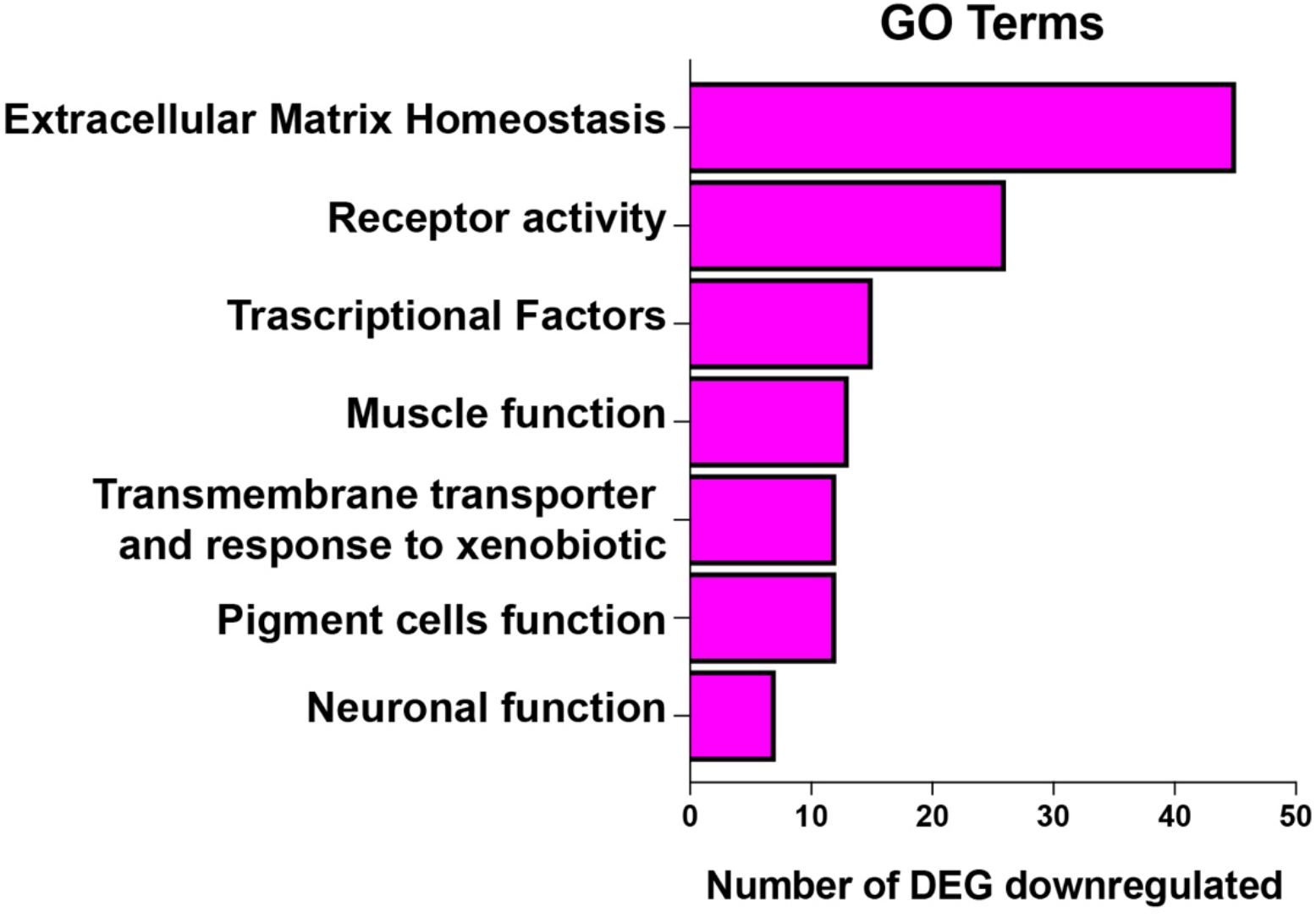
Categories of genes downregulated when FGFR is inhibited.

**Figure 7 -Figure supplement 2.**
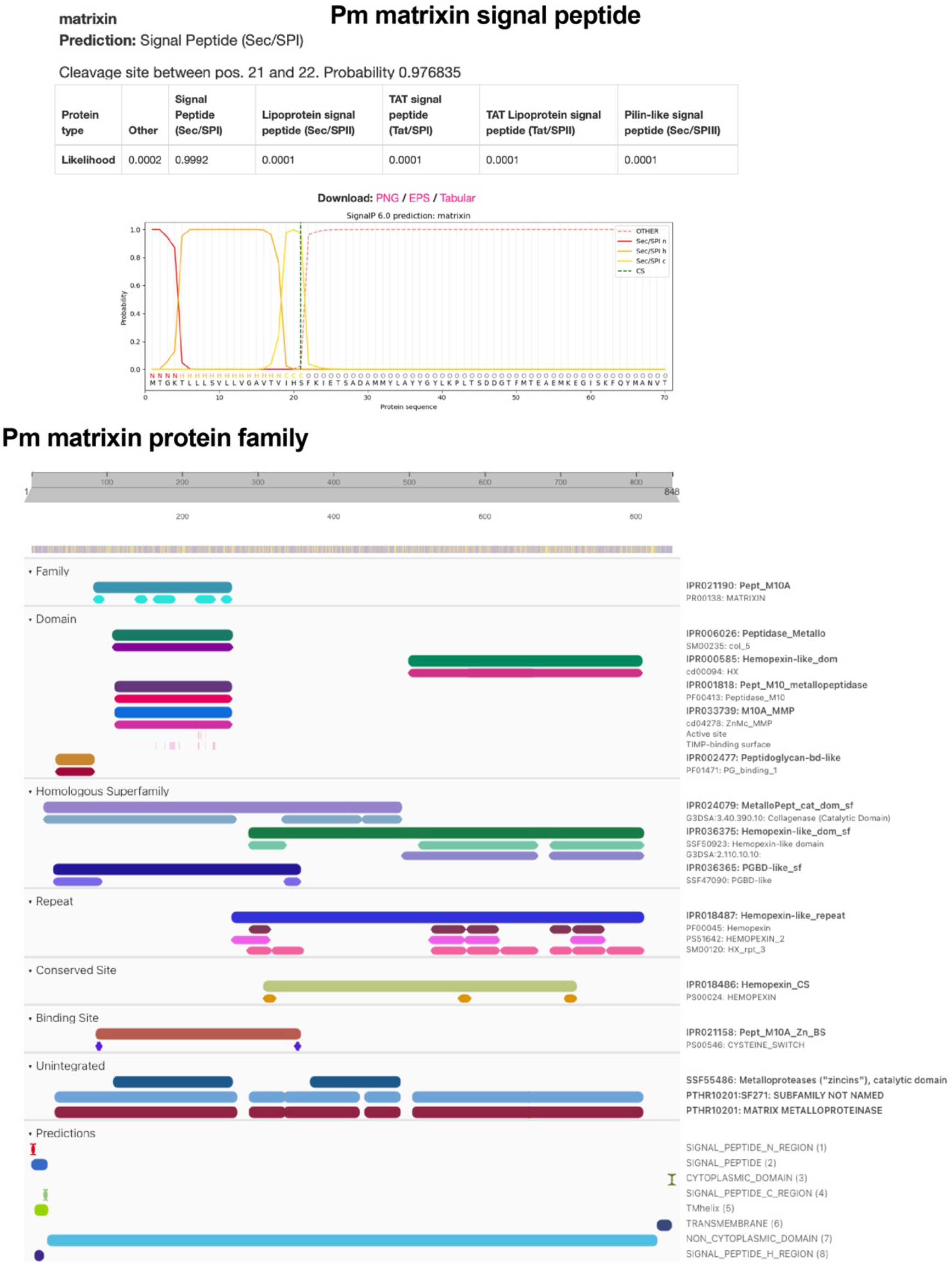
Signal peptide to identify secreted proteins and Interprot analysis of protein domain to identify protein families for matrixin.

**Figure 7 -Figure supplement 3.**
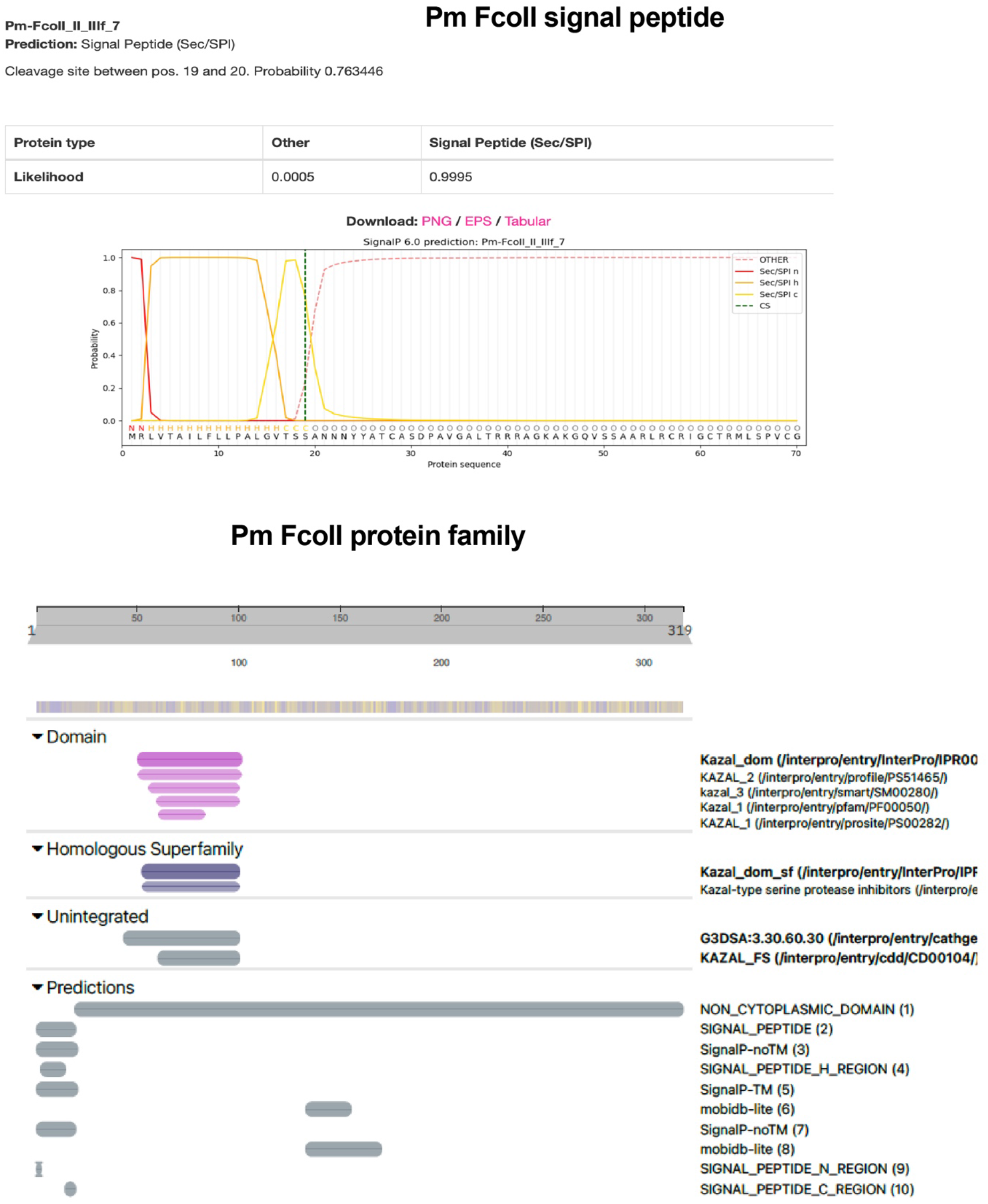
Signal peptide to identify secreted proteins and Interprot analysis of protein domain to identify protein families for FcoII.

**Figure 7- Figure supplement 4.**
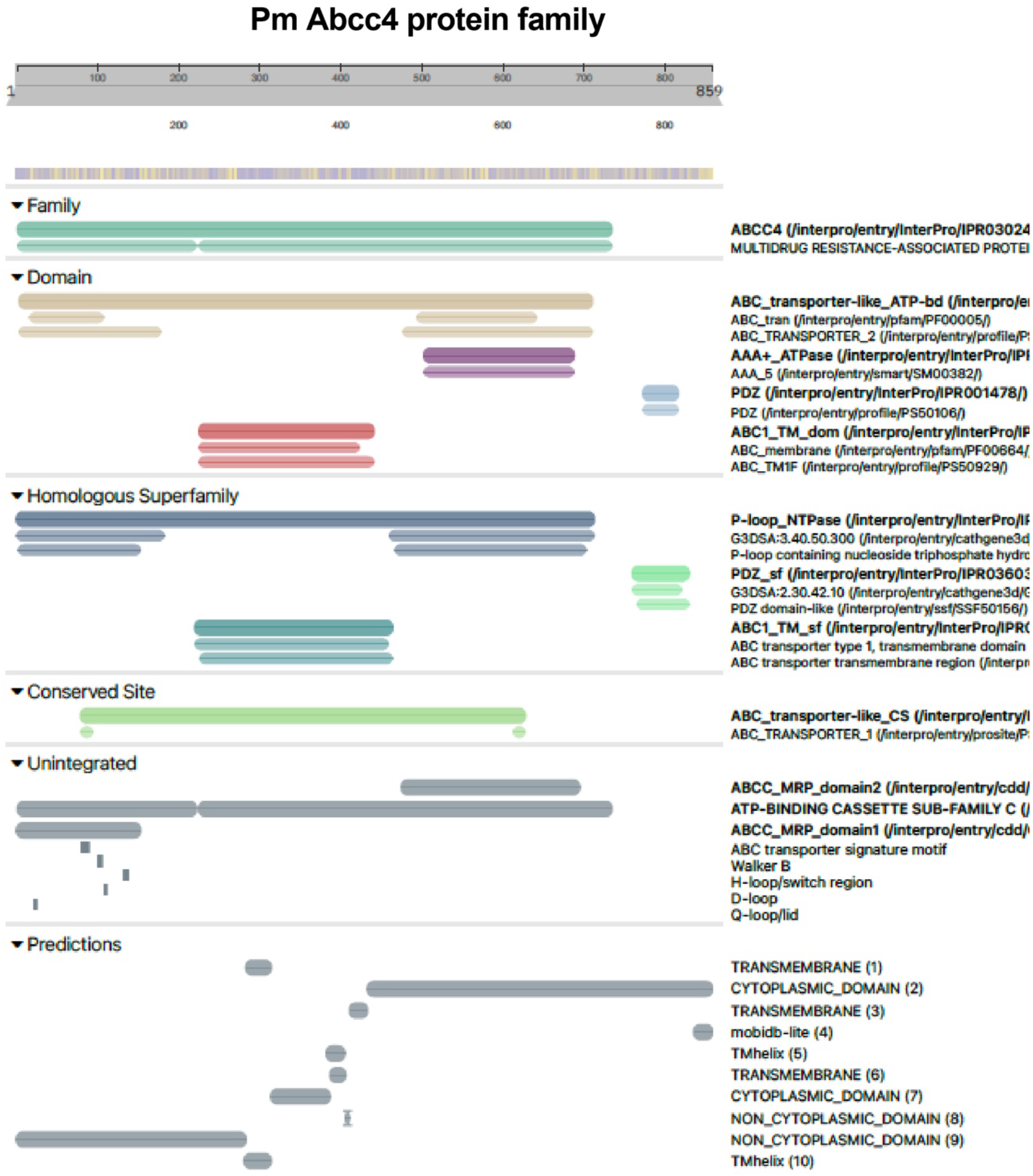
Interprot analysis of protein domain to identify protein families for the transporter Abcc4.

**Figure 7- Figure supplement 5.**
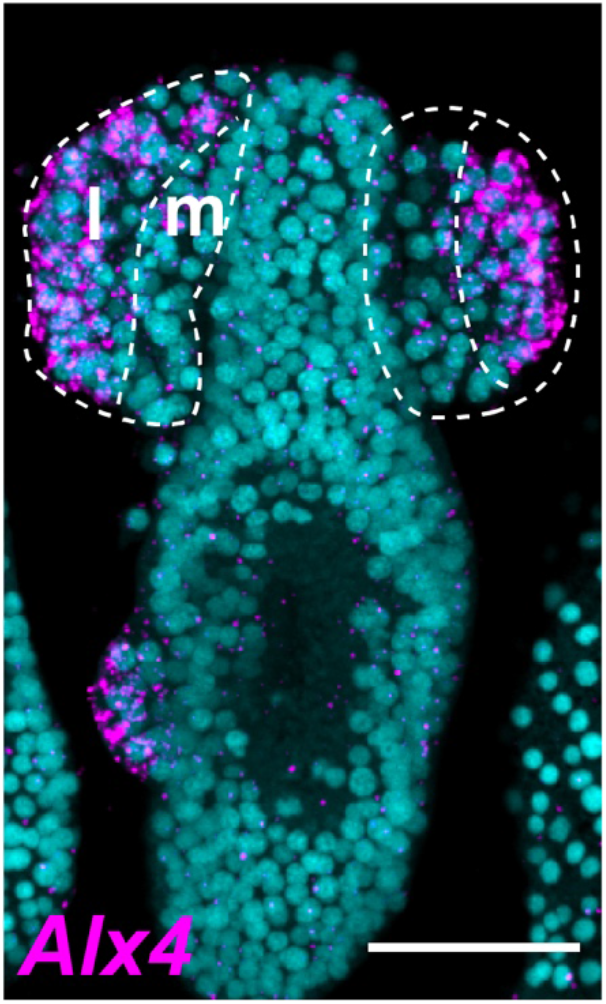
FISH for Alx4. L= lateral; m= medial. Tubes are highlighted by dotted lines. Scale bar is 50 μm.

**Figure 7- Figure supplement 6.**
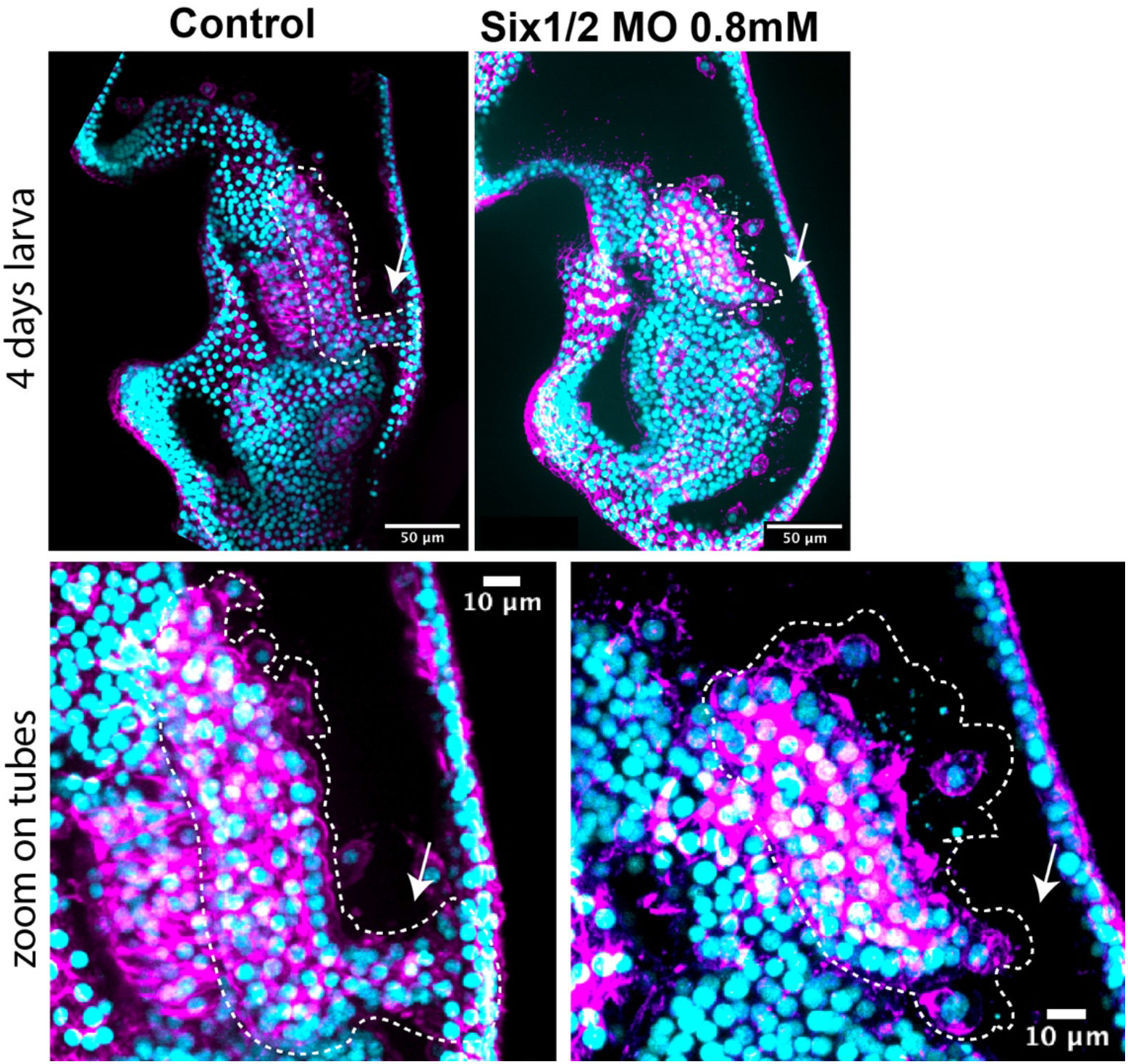
Larvae knocked down with Six1/2 morpholino showing lack of hydropore canal.

