## Supplementary material for "Molecular mechanisms of tubulogenesis revealed in the sea star hydro-vascular organ": Figure 1-Table 1

| Sea star stages | Developmental timing* | Hydro-vascular organ naming | Features |
|---|---|---|---|
| **Gastrula (G)** 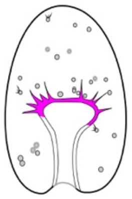 | 44-48hpf | Precursor cells (mesoderm) | Hydro-vascular organ precursor cells are located on the tip of the growing gut and are polarized. |
| **Late gastrula (LG)** 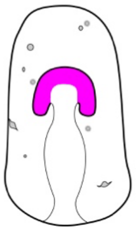 | 60-72hpf | Tubes | Tubulogenesis starts. From the tip of the gut the precursor cells start to migrate on the sides of the gut. Two tubes grow towards the posterior end of the larva. |
| **Early larva (EL)** 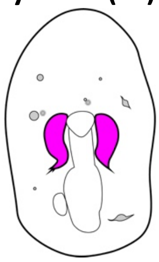 | 4d | Tubes | Two bilateral tubes are formed on the left and right sides of the gut. The two tubes detach from the gut. The hydropore canal forms. |
| **Larva (L)** 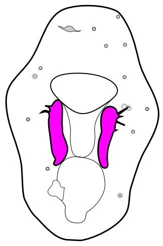 | 5-7d | Tubes | Tubes elongate. This stage lasts 3 days that we define L1 (5d), L2 (6d) and L3 (7d) each subsequent day of the larva stage. |
| **Late larva (LL)** 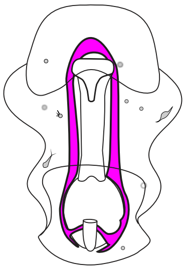 | >1w | Hydro-vascular organ | The two tubes merge to form the hydro-vascular organ, a continuous tube that has one opening towards the outside environment, the hydropore canal. |

*Hours post fertilization (hpf); Days (d); Week (w)
